# A dynamic molecular glue targets a plastic MDA5 filament interface to stimulate innate immunity

**DOI:** 10.64898/2026.10.14.759171

**Authors:** Lijuan Yan, Ziqing Ye, Zehua Li, Swechha M. Pokharel, Tiffany Hsu, Celia Torres-Arancivia, Koji Haratani, Elena V. Ivanova, Cloud P. Paweletz, Sara Cherry, David A. Barbie, Sun Hur

## Abstract

Innate immune signaling is a double-edged sword: uncontrolled activation drives autoinflammatory diseases, while controlled activation enhances antiviral and antitumor immunity. MDA5, a cytosolic dsRNA sensor that activates type I interferon signaling, mediates both immunopathology and protective immunity, yet small-molecule modulators have remained elusive. Here we report MA2, a first-in-class selective MDA5 agonist that acts as a molecular glue. MA2 binds a cryptic inter-protomer pocket within MDA5 filaments that undergoes ATPase-associated conformational remodeling. Rather than rigidly locking the interface, MA2 remains bound while accommodating and reshaping these structural transitions, strengthening MDA5–dsRNA and inter-protomer interactions and attenuating ATP hydrolysis. MA2 does not induce new filament assembly but potentiates pre-existing filaments, enabling context-dependent activation in dsRNA-enriched cells. MA2 enhances antiviral activity and synergizes with chemotherapeutic agents and natural killer cells to promote tumor cell killing. These findings establish dynamic molecular glues as a strategy for targeting conformationally plastic interfaces previously considered undruggable.

## Introduction

Therapeutic modulation of innate immune pathways that activate type I interferon (IFN) signaling is of broad interest across biology and medicine^1-3^. Pharmacological activation of the DNA sensor adaptor STING^4-7^ or inhibition of the type I IFN signaling repressor PTPN1/2^8^ synergizes with immune checkpoint therapy by inflaming the tumor microenvironment and enhancing antitumor immune responses. Similarly, therapeutic activation of the RNA sensors MDA5 and RIG-I has gained increasing attention^9-12^, particularly because, unlike cGAS-STING, which is frequently suppressed in tumors^13-15^, these sensors are more broadly expressed^16^. Despite the growing interest, however, pharmacological activation of MDA5 and RIG-I remains challenging. Many therapeutic interventions that engage MDA5 and RIG-I, including radiotherapy, DNA-damaging chemotherapy and inhibitors of chromatin and RNA biology, globally perturb cell physiology and activate these sensors indirectly by increasing endogenous dsRNA^17-26^. Several RIG-I agonists have been reported, including synthetic RNA ligands^9,10,12^ and a small molecule^27^, but the RNA-based agonists require formulation strategies to achieve stability and cellular delivery, whereas the reported small molecule has an unclear binding mode and mechanism of action. Meanwhile, MDA5 activation has largely been limited to dsRNA mimetics, such as poly(I:C), a poorly defined, heterogeneous polymer that simultaneously activates multiple dsRNA-sensing pathways including PKR and OASes/RNase L^28^. Direct and selective small-molecule activators of MDA5 have not been reported.

Previous biochemical and structural studies have shown that MDA5 functions through dynamic filament assembly^29^, which likely contributes to the challenge in developing small molecule modulators. In the resting state, MDA5 exists as a monomer, but upon binding to long dsRNA MDA5 cooperatively assembles into filaments along the length of dsRNA^30,31^. Filament assembly allows recruitment of the essential cofactor TRIM65 and activation of the signaling adaptor molecule MAVS^31-33^, triggering a signaling cascade that culminates in the induction of type I IFN as well as proinflammatory cytokines, such as CCL5 and IL6. Notably, dsRNA binding promotes MDA5’s ATP hydrolysis activity within its helicase domain, which drives conformational changes and filament destabilization^30,34,35^, providing an intrinsic regulatory mechanism to limit filament persistence and restrain MDA5 signaling (Figure 1A). Consistent with this model, mutations in the ATP active site of MDA5 often exhibit gain-of-function signaling activity in cells^29,36^.

**Figure 1.**
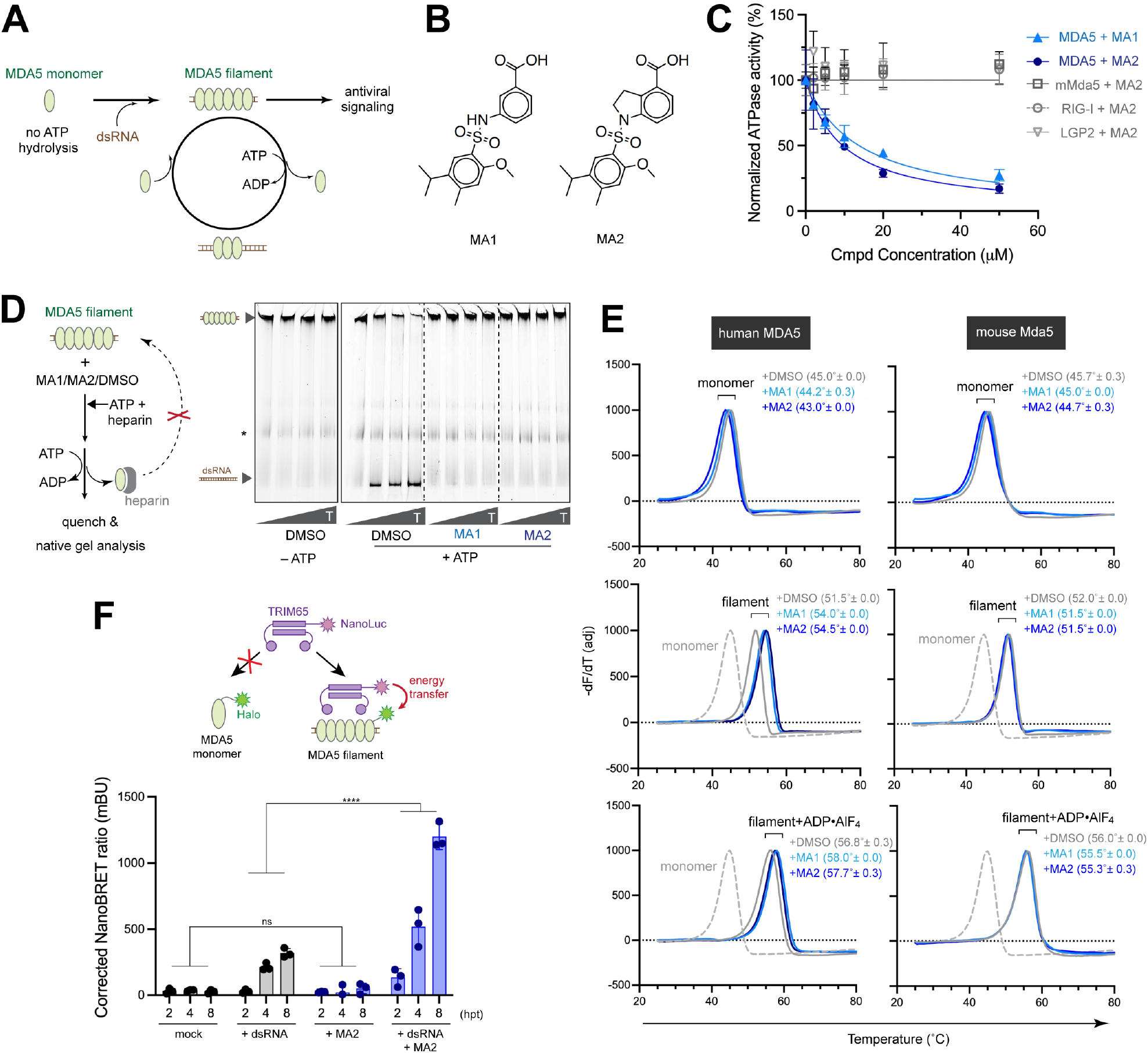
MA1 and MA2 stabilize filamentous MDA5 *in vitro* and in cells. (A) Schematic of MDA5 signaling and ATP-driven filament dynamics. MDA5 assembles into filaments upon dsRNA binding, which stimulates ATP hydrolysis within its helicase domain. ATP hydrolysis in turn promotes MDA5 dissociation from filament ends, driving disassembly. The dynamic equilibrium between filament assembly/extension and ATP-driven end-disassembly allows MDA5 to regulate filament stability according to the length of the underlying dsRNA scaffold (Ref. 35). Because only filamentous—not monomeric—MDA5 activates downstream antiviral signaling (Ref. 32, 33), this dynamic instability provides a mechanism for dsRNA length-dependent immune activation. (B) Chemical structures of MA1 and MA2. (C) Effect of MA1/MA2 on the ATPase activity of MDA5. MDA5 (200 nM) was incubated with 512 bp dsRNA (1 ng/μL) and MA1/MA2 (indicated concentration, DMSO-normalized), and reactions were initiated by ATP addition and quenched at 0.5 min. Equivalent condition was used for human RIG-I, LGP2 and mouse Mda5 (mMda5). Data are shown as average ± S.D. from three biological repeats. N-terminal CARD deletion construct was used for human MDA5 and mMda5 as they bind dsRNA and forms filaments similarly to full-length proteins, while displaying greater biochemical stability. (D) Heparin chase assay measuring MDA5 filament stability. Pre-assembled MDA5 filaments on 512 bp dsRNA were challenged with heparin (50 μg/mL) in the presence or absence of ATP (2 mM) and MA1 or MA2 (50 μM) for 0, 15, 30, 60 seconds (T). Heparin sequesters dissociated MDA5 and prevents rebinding to dsRNA, allowing filament disassembly to be monitored over time. * The bands were non-specific species from heparin. (E) Thermal shift assays measuring compound binding to monomeric or filamentous MDA5 (human MDA5 or mouse Mda5). MDA5 monomer or filament assembled on 512 bp dsRNA was incubated with MA1 or MA2 (200 μM) in the presence and absence of ADP•AlF4 (0.5 mM). The melting temperature (Tm, in parenthesis) was calculated from the inflection point of the derivative of fluorescece with respect to temperature (−dF/dT). Amplitude of the derivative was adjusted for visual clarity. Tm was derived from 3 biological repeats (mean±S.D.). (F) NanoBRET assay monitoring MDA5 filament formation in cells. Top: schematic of the assay. TRIM65 binds MDA5 specifically in its filamentous state, enabling energy transfer from TRIM65 fused to NanoLuciferase (NanoLuc) to MDA5 fused to HaloTag labeled with a Halo-618 ligand. Bottom: A549 cells stably co-expressing TRIM65-NanoLuc and MDA5-HaloTag were transfected with 512 bp dsRNA and treated with MA2, and NanoBRET signal was measured at the indicated times post-transfection. Data are shown as average ± S.D. from three biological repeats. \*\*\*\**p* <0.0001 was calculated by two-way ANOVA analysis, comparing the signal values between DMSO and MA2 treatments. ns: not significant.

Here, we report MA2, the first direct and selective small-molecule agonist of MDA5 with a unique mode of action. MA2 acts as a dynamic molecular glue that exploits the conformational plasticity of the MDA5 filament, stabilizing signaling-competent MDA5 filaments through both ATP-dependent and ATP-independent mechanisms to potentiate antiviral and anti-tumor signaling.

## Results

### MA1 and MA2 stabilize filamentous MDA5

We obtained two structurally related compounds as gifts from Odyssey Therapeutics, which had performed a small-molecule library screen using an ATPase activity assay to identify MDA5 modulators. MA1 emerged from the primary screen based on its apparent inhibition of MDA5 ATP hydrolysis, whereas MA2 was derived through subsequent structure-activity relationship optimization (Figure 1B). Our ATPase assays (measured at 0.5 min) showed that both MA1 and MA2 decreased the ATPase activity of recombinant human MDA5 in a dose-dependent manner, with apparent IC_50_ of 13.8 μM and 9.5 μM for MA1 and MA2, respectively (Figure 1C). Neither compound affected the ATPase activity of closely related paralogs RIG-I and LGP2, or mouse Mda5 (Figures 1C for MA2 and S1A for MA1). These findings indicate that MA1 and MA2 are highly specific for human MDA5. Unless otherwise specified, all MDA5 data refer to the human protein.

Given that ATP hydrolysis promotes filament disassembly, we asked whether MA1/MA2 stabilize the MDA5 filament. We monitored disassembly of pre-assembled MDA5 filament over time by capturing dissociated MDA5 proteins with excess polyanionic heparin, which prevented their rebinding to dsRNA^30^ (Figure 1D, left). Consistent with previous reports^30,34^, ATP promoted rapid filament disassembly in the absence of MA1/MA2 (Figure 1D). In contrast, MA1/MA2 largely preserved filament integrity, maintaining filament stability at levels similar to those observed in the absence of ATP (Figure 1D). These results demonstrate that MA1/MA2 counteract ATP-driven filament disassembly.

To determine whether MA1 and MA2 directly bind MDA5, we performed thermal shift assays, which are based on the principle that specific ligand binding often increases thermal stability of a target protein^37^. As expected, filament assembly of MDA5 on 512 bp dsRNA increased Tm from 45.0°C for a monomeric MDA5 to 51.5°C for filamentous MDA5 (Figure 1E, left). ADP•AlF_4_, an ATP hydrolysis transition state mimic, further stabilized MDA5 filament, increasing Tm to 56.8°C (Figure 1E, left). Neither MA1 nor MA2 stabilized monomeric MDA5, but instead decreased monomeric MDA5 Tm by ∼1-2°C (Figure 1E, top left). In contrast, both compounds stabilized filamentous MDA5, both in the presence and absence of ADP•AlF_4_ (Figure 1E, center and bottom left). No stabilization was observed for mouse Mda5 filament (Figure 1E, right) or dsRNA-bound RIG-I or LGP2 (Figure S1B), although modest destabilization of monomeric proteins was observed across all proteins tested. These results suggest that MA1/MA2 selectively bind and stabilize the filamentous form of human MDA5 without competing with ATP binding.

To further test the impact of MA2 on filament stability in cells, we developed a NanoBRET assay^38^ to monitor proximity between MDA5 and its functional partner TRIM65, which binds MDA5 specifically in its filamentous form and activates MDA5 signaling^32^. In this system, TRIM65 was fused to Nano-Luciferase (NanoLuc) and MDA5 was tagged with Halo protein, allowing energy transfer from NanoLuc to Halo-conjugated 618 ligand upon their interaction (Figure 1F, top). This assay therefore provides a proxy for cellular MDA5 filament formation. Consistent with the requirement for dsRNA in MDA5 filament formation and TRIM65 binding, dsRNA transfection increased NanoBRET signals, and MA2 treatment further enhanced these signals only in the presence of dsRNA (Figure 1F, bottom), again consistent with MA2 selectively acting on filamentous MDA5.

Together, these findings demonstrate that MA1 and MA2 selectively engage and stabilize the filamentous, signaling-competent state of MDA5 *in vitro* and in cells.

### MA1 and MA2 stimulate MDA5 signaling

Given that filament assembly is required for MDA5 signaling, we next examined how MA1/MA2 stimulate the pathway in cells. We used the breast cancer cell MCF-7 and the non-small cell lung cancer cell HCC2429, which have an intact endogenous MDA5 signaling pathway. Both compounds stimulated antiviral signaling, as measured by IFNβ ELISA (Figure 2A), with MA2 displaying substantially greater potency despite their similar *in vitro* activity, likely reflecting improved cell permeability. We therefore focused subsequent analyses on MA2 and further confirmed its immune stimulatory activity by measuring STAT1 phosphorylation, *IFNB* mRNA and CXCL10 secretion, all of which were dependent on endogenous MDA5 (Figures 2B, 2C, and S2A).

**Figure 2.**
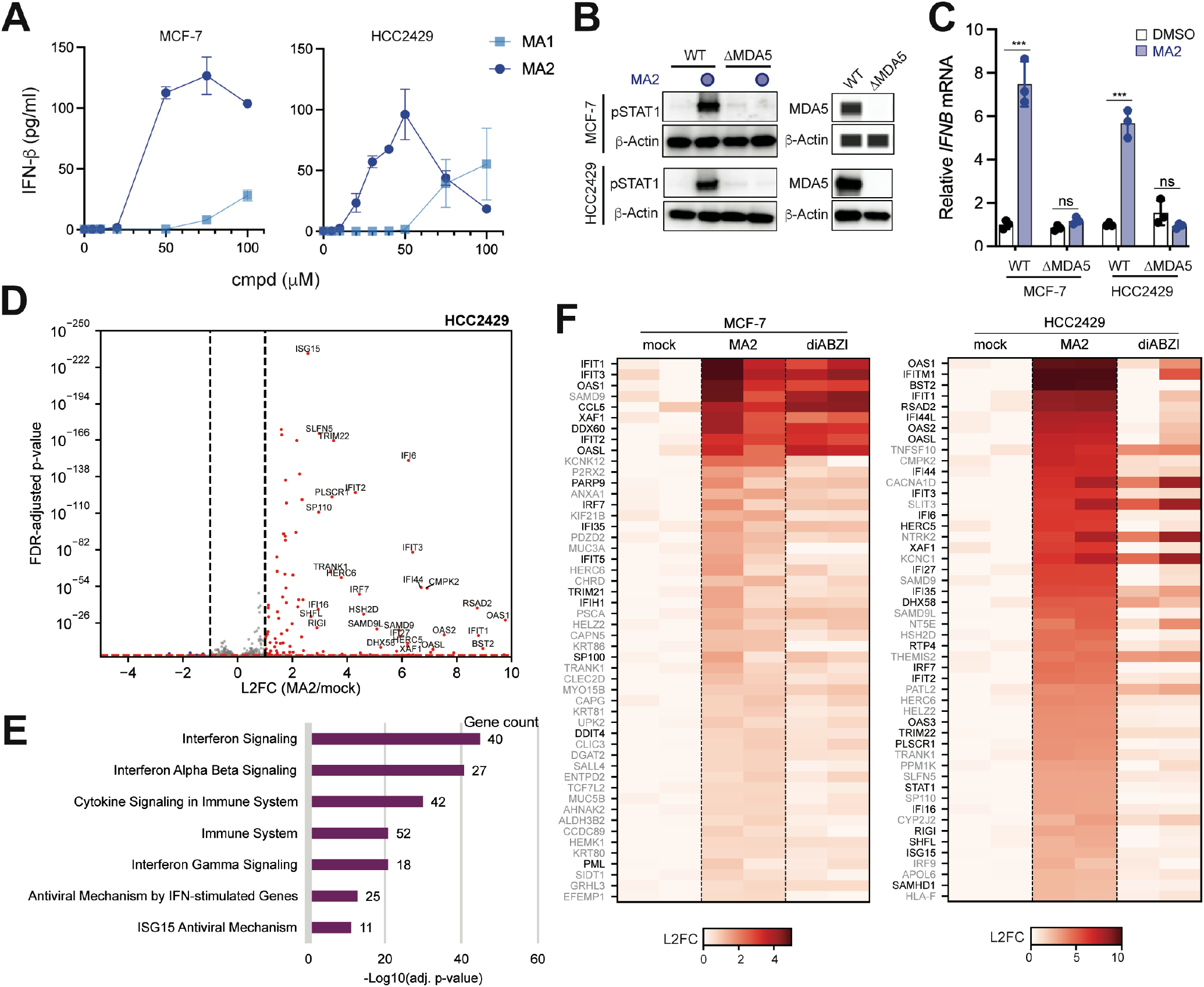
MA1 and MA2 activate MDA5 signaling. (A) Dose-dependent induction of IFNβ by MA1 and MA2 in MCF-7 and HCC2429 cells, as measured by IFNβ ELISA. Data are shown as average ± S.D. from three biological repeats. (B) Immunoblot analysis of STAT1 phosphorylation (pSTAT1) in wild-type (WT) and MDA5 knockout (ΔMDA5) MCF-7 or HCC2429 cells treated with MA2 (50 μM). Jess Automated Western System was used for analyzing MDA5 and β-actin in MCF-7. (C) RT-qPCR analysis of *IFNB* mRNA induction in response to MA2 (50 μM) in WT and ΔMDA5 cells. Data are shown as average ± S.D. from three biological repeats. \*\*\**p* <0.001 was calculated by two-tailed student’s t-test, ns: not significant. (D) Volcano plot of differentially expressed genes in HCC2429 cells upon MA2 stimulation. Significantly upregulated genes (L2FC > 1, padj < 0.05) are colored red; significantly downregulated genes (L2FC < -1, padj < 0.05) are in blue, and the rest are in grey. The names of genes with (L2FC > 2.5, padj < 0.00001) are displayed. STAT1, OAS3, HELZ2 are also upregulated (L2FC > 1) but are not shown because padj < 10-250. (E)Reactome pathway enrichment analysis of MA2-upregulated genes in HCC2429 cells (padj < 0.05, L2FC > 1). The top 7 most significantly enriched categories are shown. Adjusted p-values for pathway enrichment are calculated by EnrichR. (F)Heatmap of gene expression changes upon stimulation with MA2 (50 μM) or diABZI (10 μM). Top 50 MA2-induced genes (padj < 0.05) are shown. Interferon stimulated genes (ISGs) are in black. Note that diABZI showed toxicity above 10 μM, limiting its use at higher concentrations.

To comprehensively characterize the transcriptional consequences of MA2 treatment, we performed RNA-seq analysis. MA2 broadly induced innate immune genes, with the top 7 enriched pathways in both MCF-7 and HCC2429 cells related to antiviral and interferon signaling pathways (Figures 2D, 2E, S2B and S2C), indicating a high degree of pathway selectivity. Heatmap analysis of top MA2-induced genes revealed a transcriptional program highly similar to that induced by the STING agonist diABZI^7^ and dsRNA mimetic polyIC in MCF-7 cells (Figures 2F, left, and S2D-S2F). Interestingly, in HCC2429 cells, diABZI failed to induce a comparable antiviral transcriptional response at all doses tested (Figures 2F, right, S2G and S2H). Instead, diABZI repressed expression of histone genes, a response not observed with MA2 (Figures 2D and S2G). RT-qPCR analysis of *IFNB* expression further confirmed robust responsiveness to MA2 in both cell lines, whereas diABZI was active only in MCF-7 cells (Figure S2H). These findings demonstrate that MA2 selectively activates endogenous MDA5 signaling and can remain effective in cancer cells that are refractory to STING agonists.

### MA1 and MA2 bind a cryptic pocket formed at the MDA5 filament interface

To elucidate how MA1 and MA2 selectively recognize MDA5 filaments, we determined the cryo-EM structures of MDA5 filaments bound to each compound. For these studies, we employed the gain-of-function MDA5 variant G495R^36^, which displays enhanced dsRNA binding with improved conformational homogeneity, facilitating cryo-EM analysis. Similar to WT MDA5, the G495R variant selectively bound MA1 and MA2 in its filamentous state and displayed enhanced signaling activity upon MA2 addition (Figures S1C and S1D).

We determined three structures of MDA5 filament assembled on 512 bp dsRNA: without any compound and in complex with MA2 or MA1 at resolutions of 3.8 Å, 2.8 Å and 3.6 Å, respectively (Figures S3-S5, and Table S3). In MA2 or MA1-bound structures, density that could not be accounted for by MDA5 or dsRNA appeared in the same location within the interface between adjacent protomers (Figures 3A and 3B). Both compounds could be unambiguously fitted to the density in a single orientation. The inter-protomer location of MA2 and MA1 explains their selective interaction with filamentous but not monomeric MDA5. Given the higher-resolution reconstruction and greater cellular potency of MA2, subsequent analyses focused primarily on MA2-bound MDA5 structures.

**Figure 3.**
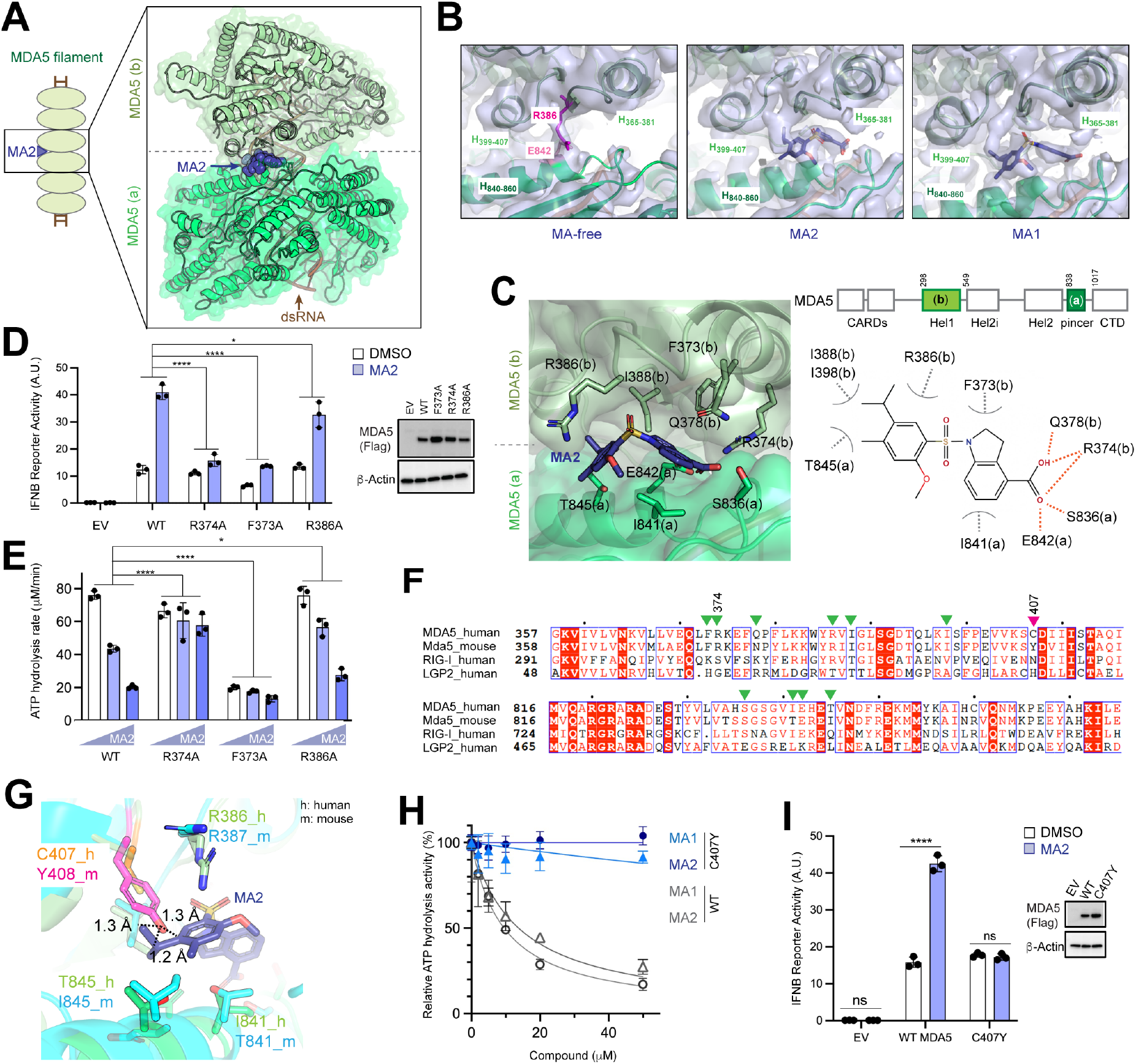
MA1 and MA2 bind a cryptic pocket formed at the MDA5 filament interface. (A) Overall structure of the MDA5 filament bound to MA2. Dimeric segments (protomers a and b) were computationally extracted from filaments for reconstruction and refinement. MDA5 protomers (green) and dsRNA (brown) are shown in cartoon with transparent surface for MDA5 proteins. MA2 is shown as spheres (dark blue). (B) Electron density at the MA-binding pocket for MA-free (left), MA2-bound (center) and MA1-bound (right) MDA5 filaments. MA2 and MA1 could be unambiguously modeled into the density in the respective bound structures. In the MA-free structure, only partial density was observed at this site, likely corresponding to R386(b) and E842(a). (C) Interaction between MA2 and surrounding residues in MDA5 protomers (a) and (b). Top right: domain architecture of MDA5. MA2-interacting residues from protomer (a) reside in the pincer domain, whereas those from protomer (b) are in the Hel1 domain. (D) Validation of MA2-interacting residues by IFNβ reporter dual luciferase assay. 293T cells lacking ADAR1 were transiently transfected with plasmids expressing FLAG-tagged MDA5 (WT or indicated mutant) and incubated with MA2 (50 μM) or DMSO. EV, empty vector control. Data are shown as average ± S.D. from three biological repeats. \*\*\*\**p* <0.0001, \**p* <0.05 were calculated by two-way ANOVA analysis, comparing MA2-induced signaling activation between WT and mutant MDA5 constructs. (E) Validation of MA2-interacting residues by ATPase assay. Experiment was performed as in Figure 1C with increasing concentrations of MA2 (0, 10, 50 μM). Data are shown as average ± S.D. from three biological repeats. \*\*\*\**p* <0.0001, \**p* <0.05 were calculated by two-way ANOVA analysis, comparing MA2-induced ATPase inhibition between WT and mutant MDA5 proteins. (F) Sequence alignment of human MDA5, RIG-I and LGP2 and mouse Mda5, focusing on MA2-interacting residues (green triangles) and adjacent positions. Pink arrow indicates C407 in human MDA5, discussed below. (G) Superposition of mouse Mda5 structure (light blue, PDB:6H61) onto human MDA5 (light green) in complex with MA2. Tyr408 in mouse Mda5 (magenta) is equivalent to Cys407 in human MDA5 (orange) and would clash with MA2 if the compound were to bind mouse Mda5 in the same orientation as in human MDA5. (H) Effect of the C407Y mutation on MA2 binding, as measured by ATPase assay (performed as in Figure 1C). Data are shown as average ± S.D. from three biological repeats. (I) Effect of MA2 on MDA5 (WT or C407Y) signaling activity using IFNβ reporter dual luciferase assay (performed as in Figure 3D). Data are shown as average ± S.D. from three biological repeats. \*\*\*\**p* <0.0001 was calculated by two-tailed student’s t test.

MA2 occupied a composite binding pocket formed by Ser836, Ile841, Glu842 and Thr845 of the pincer domain from one protomer (a) and Phe373, Arg374, Gln378, Arg386, Ile388 and Ile398 of the Hel1 domain from the neighboring protomer (b) (Figure 3C). Arg374(b), Gln378(b), Glu842(a) and Ser836(a) together interacted with the carboxylate substituent on the MA2 indoline ring through electrostatic and polar interactions, while the remaining pocket residues–Phe373(b), Ile388(b), Ile398(b) and Ile841(a)– formed a hydrophobic cradle around the indoline and isopropyl phenyl ring. The Arg386(b) side chain also nestled above the phenyl group, potentially contributing a cation-π interaction.

To validate the binding modes of MA2, we performed mutagenesis analysis. Mutation of Arg374 to Ala had little impact on the dsRNA-dependent cellular signaling activity (Figure 3D) or ATP hydrolysis activity (Figure 3E) in the absence of MA2, indicating that neither dsRNA binding nor filament formation is significantly affected by this mutation. However, the R374A mutation nearly abolished MA2’s effect on cellular signaling (Figure 3D) and ATP hydrolysis (Figure 3E), demonstrating Arg374 as a critical residue in MA2 binding. Consistent with this notion, MA2 was less efficient in stabilizing R374A filament than WT MDA5 filament (Figure S6A). Mutation of Phe373 to Ala diminished both basal signaling activity and ATP hydrolysis, as well as the impact of MA2 (Figures 3D and 3E), suggesting that Phe373 contributes to both filament integrity and the MA2 binding interface. In contrast, mutation of Arg386 to Ala had a modest effect on both basal activities and response to MA2, suggesting that Arg386 plays a more modest role in MA2 binding.

The MA2-bound MDA5 structures also explain its high specificity for human MDA5 relative to RIG-I, LGP2 and mouse Mda5. Sequence alignments showed that MA2-interacting residues, in particular Arg374 critical for MA2 binding, are not conserved in RIG-I or LGP2 (Figure 3F), accounting for the lack of compound activity towards these proteins. In contrast, these residues were largely conserved in mouse Mda5 (Figure 3F), suggesting that additional structural features determine species selectivity. We noticed that Cys407 in human MDA5, located immediately adjacent to the MA2-binding pocket, is replaced by Tyr408 in mouse Mda5, which may introduce a steric clash with MA2 (Figure 3G). Consistent with this idea, C407Y mutation in human MDA5 abolished MA2 binding, as measured by ATPase activity, thermal shift assay, and cellular signaling activity (Figures 3H, 3I, and S6B). Y408C mutation in mouse Mda5, however, did not restore MA2 binding (Figure S6C), suggesting additional contributions from other residues.

Together, these structural and functional analyses explain why MA2 selectively recognizes filamentous human MDA5, while remaining inactive toward monomeric MDA5, the related paralogs RIG-I and LGP2 and the mouse ortholog.

### MA2 accommodates ATP-driven remodeling of the filament interface

ATP binding and hydrolysis drive extensive conformational remodeling of the MDA5 filament interface^35^, including the cryptic pocket occupied by MA2. This raised the question of how MA2 accommodates ATP-driven remodeling of its binding site and, conversely, how MA2 influences ATP-dependent conformational changes of the filament. Although MA2 attenuates ATP hydrolysis, it does not abolish it (Figures S6D-S6F), indicating that ATP turnover continues to occur within MA2-containing filaments. Additionally, MA2 also bound ADP•AlF_4_-bound MDA5 filament (Figure 1E), suggesting that MA2 is compatible with ATP binding and/or hydrolysis. These observations suggest two possible scenarios: MA2 may act as a rigid molecular glue that locks the filament interface and prevents ATP-dependent conformational changes; alternatively, MA2 may dynamically accommodate ATP-driven conformational changes in the filament interface.

To distinguish between these models, we determined an additional cryo-EM structure of MDA5 filament in the presence of both MA2 and ADP•AlF_4_ (Figure S7, and Table S3). The quaternary complex structure showed that MA2 occupies the same inter-protomer pocket in the presence of ADP•AlF_4_ as in the nucleotide-free state (Figure 4A, left). Comparison of the MA2-bound filament in the presence and absence of ADP•AlF_4_ revealed distinct filament geometries that differed by ∼5° in helical twist and 1.2 Å in rise (Figure 4A, left, and 4B). Although ADP•AlF_4_ also altered filament geometry in the absence of MA2 (Figure 4A, right), ADP•AlF_4_-induced changes in the presence of MA2 differed substantially from those observed in MA2-free filaments, which was also reflected in distinct helical parameters (Figure 4B).

**Figure 4.**
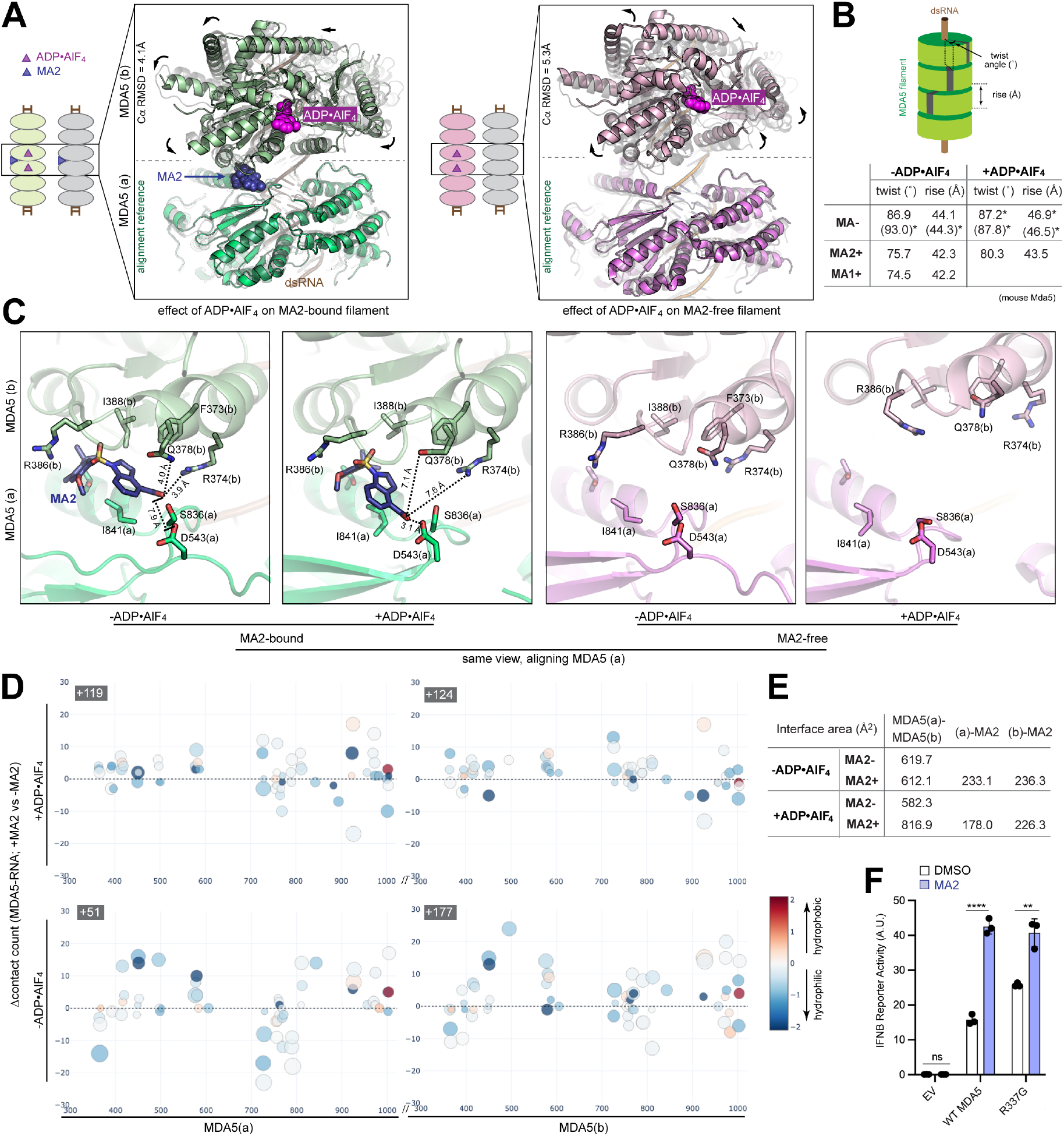
MA2 remains bound during ATP-driven filament remodeling, while enhancing MDA5-RNA and inter-protomer interactions. (A) Conformational changes induced by ADP•AlF4 in MA2-bound (left) and MA2-free (right) MDA5 filaments. Nucleotide-free structures (transparent grey) were superimposed onto the corresponding ADP•AlF4-bound dimeric segments of MA2-bound (green) or MA2-free (pink) MDA5 filaments. Ca RMSD was calculated for protomer (b), while aligning on protomer (a). Black arrows indicate the direction of movement upon ADP•AlF4 binding. (B) Effect of MA2, MA1 and ADP•AlF4 on MDA5 filament helical geometry. Helical twist and rise parameters for MDA5 filaments in the apo and ADP•AlF4 states, with or without MA1/MA2 binding were obtained by Chimera. *Values are derived from previously published structures (PDB:7JL2, 6H66, 6GKH). Values in parenthesis are from mouse MDA5 filament structures. (C) Close-up view of the MA2 pocket and its remodeling upon ADP•AlF4 binding in MDA5 filament, with (left two) and without (right two) MA2. All views are from the same orientation, aligned on protomer (a). (D) Effect of MA2 on MDA5-RNA interaction. Each point represents a net change in the contact count for a given protein residue in subunit (a) or (b) summed over all RNA residues. Positive values indicate a net gain of RNA contact upon MA2 binding, while negative values indicate a net loss. Contacts were defined as heavy-atom pairs within 4.5 Å. The net contact count change summed over all protein residues for a given protomer is shown in grey box. Dot size represents total RNA contact count in whichever of the two structures (+MA2 or −MA2) shows the stronger interaction for that residue; dot color represents hydrophobicity/hydrophilicity of that residue according to the Wimley-White scale. (E) Interface areas (Å2) of the MDA5 filament structures were calculated using PDBePISA, including the inter-protomer (MDA5(a)–MDA5(b)) interface and the compound-protein interfaces. (F) Effect of MA2 on MDA5 (WT or R337G) signaling activity using IFNβ reporter dual luciferase assay (performed as in Figure 3D). Data are shown as average ± S.D. from three biological repeats. \*\*\*\**p* <0.0001, \*\**p* <0.01 were calculated by two-tailed student’s t test.

These global changes in the filament architecture resulted in local remodeling of the MA2-binding pocket and its interaction with MA2 (Figure 4C). Arg374(b) and Gln378(b), which contacted MA2 carboxylate in the nucleotide-free state, no longer interacted with the compound in the presence of ADP•AlF_4_. Conversely, Asp543(a) from the adjacent protomer approached MA2 carboxylate upon ADP•AlF_4_ binding, reducing the distance from 7.9 Å to 3.1 Å, which can support a water-mediated interaction. Other residues, such as Arg386(b) and Ser836(a), moved along with MA2 upon ADP•AlF_4_ binding to adapt to changes in MA2-MDA5 interaction. Thus, rather than recognizing a rigid protein interface, MA2 accommodates ATP-dependent filament remodeling through a flexible interaction network. Importantly, these ATP-dependent conformational changes in the MA2-binding pocket differed markedly from those observed in the absence of MA2 (Figure 4C), indicating that MA2 not only tolerates filament remodeling during ATP hydrolysis, but also actively reshapes the structural response of the filament to ATP binding and hydrolysis.

### MA2 strengthens MDA5-RNA and inter-protomer interactions to stabilize filaments in ATP-dependent and -independent mechanisms

We next asked how MA2 stabilizes the MDA5 filament and whether MA2-induced conformational changes can explain MA2’s effect on ATP hydrolysis and filament stability. Superposition of MDA5–ADP•AlF_4_ structures in the presence and absence of MA2 showed that both the catalytic residues surrounding ADP•AlF_4_ and overall conformation of individual MDA5 protomers remained nearly identical (Figure S6G), arguing against a direct effect of MA2 on the ATP-binding pocket. In contrast, MA2 markedly altered filament architecture. In both nucleotide-free and ADP•AlF_4_-bound states, MA2 induced distinct filament conformations characterized by reduced helical twist (Δtwist=11°or 7°, respectively) and rise (Δrise=1.8 Å or 3.4 Å, respectively) relative to corresponding MA2-free structures (Figure 4B), resulting in a ∼11 Å shift in one subunit relative to the adjacent subunit (Figure S6H). Similar changes were introduced by MA1, suggesting a shared mechanism of action (Figures 4B and S6I). These observations thus suggest that MA2 attenuates ATP hydrolysis, not by directly affecting the ATP binding pocket, but by remodeling the filament geometry that is coupled to ATPase activity.

We next examined how this remodeling influences molecular interactions within the filament. Analysis of protein-RNA contacts revealed globally increased interactions between MDA5 and dsRNA throughout the central RNA-binding channel in both the nucleotide-free and ADP•AlF_4_-bound states (Figure 4D). Although the direct protein-protein interface between neighboring protomers changed little in the nucleotide-free state, it increased in the presence of ADP•AlF_4_ (Figure 4E). Moreover, MA2 forms extensive contacts with both adjacent protomers, substantially increasing the total buried solvent accessible surface area (i.e. 2X interface) in both nucleotide states (Figure 4E). Thus, MA2-induced filament remodeling strengthens MDA5-dsRNA and inter-protomer contacts.

These observations also suggested that MA2 may stabilize MDA5 filaments independently of its attenuation of ATP hydrolysis. To test this, we examined the impact of MA2 on the ATPase-deficient gain-of-function mutant R337G^36^. Despite its defective ATP hydrolysis, R337G signaling was further enhanced by MA2 (Figure 4F), indicating that MA2 can potentiate MDA5 signaling through a mechanism independent of ATP hydrolysis. This finding is consistent with the observation that MA2 enhanced filament thermal stability even in the absence of ATP hydrolysis (Figure 1E). Together, these results support a model in which MA2 stabilizes MDA5 filaments through two complementary mechanisms: (i) attenuating ATP hydrolysis and the associated filament turnover, and (ii) remodeling filament geometry to strengthen MDA5-dsRNA and inter-protomer interactions.

### MA2 elicits antiviral and antitumor immunity and synergizes with dsRNA-inducing cancer therapies

To assess the therapeutic potential of MA2, we first examined whether MA2-mediated innate immune activation confers antiviral protection. We challenged MCF-7 and HCC2429 cells with encephalomyocarditis virus (EMCV) at MOIs of 0.01 and 0.03, respectively, and viral RNA was quantified at 24 hours post-infection. MA2 potently reduced the level of EMCV RNA in both cell types, and this antiviral activity was dependent on MDA5 (Figure 5A). MA2 similarly restricted SARS-CoV-2 replication in Calu-3 cells, as measured by immunofluorescent detection of the viral spike protein (Figure 5B). In contrast, MA2 had no effect on influenza A virus (IAV) (Figure 5C), consistent with the established role of MDA5 in sensing picornaviruses and coronaviruses but not orthomyxoviruses^39,40^. These results further confirm the MDA5 specificity of MA2.

**Figure 5.**
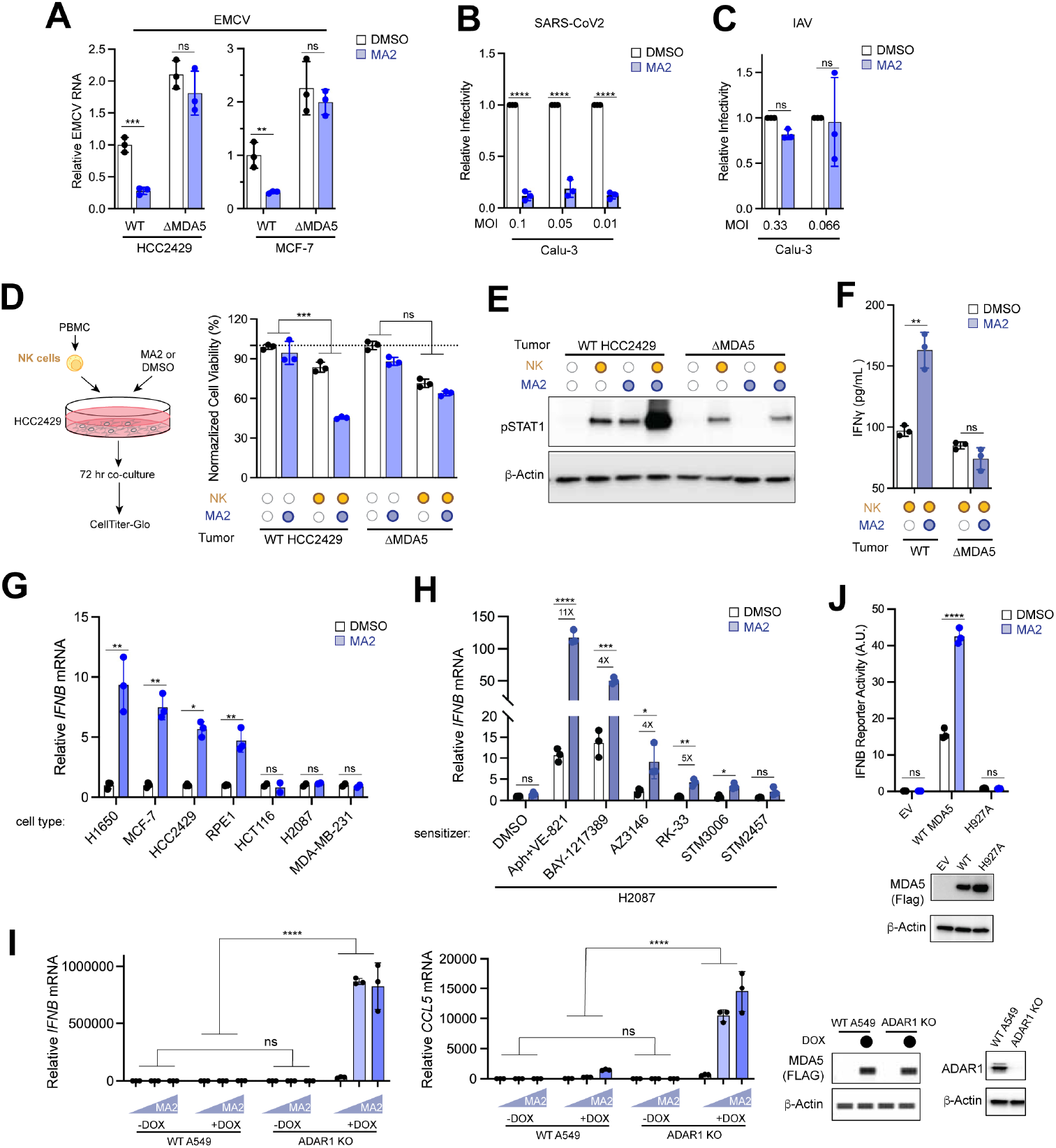
MA2 elicits antiviral and antitumor immunity and synergizes with dsRNA-inducing cancer therapies. (A) Effect of MA2 on EMCV replication. HCC2429 and MCF-7 cells were pretreated with or without MA2 (50 μM), infected with EMCV (MOI = 0.03 for HCC2429 and MOI = 0.01 for MCF-7), and harvested 24 h post-infection. EMCV RNA was quantified by RT-qPCR with GAPDH as an internal control and plotted relative to mock-infected, DMSO-treated controls. \*\*\**p* <0.001, \*\**p* <0.01 were calculated by two-tailed student’s t test. (B-C) Effect of MA2 on SARS-CoV2 (B) or IAV replication (C). Calu-3 cells were pre-treated with MA2 (50 μM) and infected with SARS-CoV-2 and IAV at indicated MOI. Cells were harvested 48 hrs post-infection and viral replication was measured by immunofluorescent analysis of SARS-CoV-2 spike protein or IAV proteins. \*\*\*\**p* <0.0001, was calculated by two-tailed student’s t test. (D)Effect of MA2 on NK cell-mediated tumor killing. HCC2429 (target) cells were treated with MA2 (50 μM) and co-cultured with primary human NK cells at an NK-to-target ratio of 1:5 for 72 hrs. HCC2429 cell viability was measured by CellTiter-Glo luminescence, with the NK cell signal subtracted and values normalized to DMSO-treated controls. \*\*\**p* <0.001 was calculated by two-way ANOVA analysis, comparing the effect of MA2 on tumor cell killing in the presence versus absence of NK cells, assessing the interaction between MA2 and NK-cell. (E)Effect of MA2 on STAT1 phosphorylation in tumor-NK cell co-culture. HCC2429 cells were co-cultured with NK cells at an NK-to-target ratio of 1:10 with or without MA2 (50 μM). After 48 hrs of co-culture, NK cells were removed by PBS washing, and HCC2429 cell lysates were collected for immunoblot analysis. (F) Effect of MA2 on IFNγ expression in tumor-NK co-culture. HCC2429 cells were co-cultured with NK cells for 48 hrs as in (d) and secreted IFNγ was measured by ELISA. \*\*\**p* <0.001, was calculated by two-tailed student’s t test. (G) RT-qPCR analysis of *IFNB* induction in response to MA2. Indicated cell lines were incubated with MA2 (50 μM) and harvested 24 hr later for analysis. *IFNB* mRNA level was internally normalized against *GAPDH* mRNA. \*\**p* <0.01, \**p* <0.05 were calculated by two-tailed student’s t test. (H)MA2-unresponsive cell H2087 was pre-treated with indicated compounds (0.5 μM aphidicolin + 2.5 μM VE-821, 200 nM BAY-1217389, 1 μM AZ3146, 0.3 μM RK-33, 0.5 μM STM3006, 1 μM STM2457) for 24 hrs, followed by treatment with MA2 (50 μM) for additional 24 hrs prior to harvest. \*\*\*\**p* <0.0001, \*\*\**p* <0.001, \*\**p* <0.01, \**p* <0.05 were calculated by two-tailed student’s t test. (I) Induction of *IFNB* or *CCL5* mRNA expression in WT A549 cells or ADAR1-knockout cells with doxycycline-inducible MDA5 expression. Cells were treated with increasing concentrations of MA2 (0, 20, 50 μM) with or without doxycycline (DOX). Jess Automated Western System was used for MDA5 and β-actin immunoblotting. \*\*\*\**p* <0.0001 was calculated by two-way ANOVA analysis, comparing the effect of MA2 on type I IFN signaling activation in the presence versus absence of ADAR1, reflecting the role of dsRNA accumulation in MA2 response. (J)Effect of MA2 on MDA5 (WT or H927A) signaling activity using IFNβ reporter dual luciferase assay (performed as in Figure 3D). \*\*\*\**p* <0.0001 was calculated by two-tailed student’s t test. All data are shown as average ± S.D. from three biological repeats.

We next investigated whether MA2-mediated innate immune activation could be harnessed for antitumor immunity. Given that MA2 robustly induced proinflammatory cytokines and chemokines (Figure S8A) without causing detectable cellular toxicity, we asked whether MA2 could synergize with immune cells, such as natural killer (NK) cells, for tumor cell killing. Because MA2 is specific to human MDA5, and does not act on mouse Mda5 (Figures 1C-1E), we focused on *in vitro* co-culture assays examining the impact of NK cells on human tumor cell lines. MA2 significantly enhanced NK cell-mediated killing of MCF-7 and HCC2429 cells in a tumor-intrinsic MDA5-dependent manner (Figures 5D and S8B). MA2 alone without NK cells had minimal effects on tumor cell viability (Figures 5D and S8B), indicating that its antitumor activity is mediated primarily through NK cells rather than from direct cytotoxicity.

We next asked how MA2 enhances NK cell-mediated tumor killing. Analysis of IFNγ––the signature NK effector cytokine––and STAT1 phosphorylation revealed that both increased upon addition of MA2 in tumor-NK cell co-culture, in a manner dependent on tumor-intrinsic MDA5 (Figures 5E, 5F, S8C and S8D). Such enhancement of IFNγ–STAT1 signaling was significantly less efficient when NK cells or tumor cells were cultured separately with MA2 (Figures 5E, and S8C-S8E). Thus, MA2’s action on tumor cells amplifies reciprocal signaling with NK cells, providing a potential mechanism for enhanced NK-mediated tumor killing.

To assess how broadly MA2 activates MDA5 signaling across cell types, we profiled seven human cell lines as measured by *IFNB* induction. Four of these lines, H1650 (non-small cell lung cancer), MCF-7 (breast cancer), HCC2429 (non-small cell lung cancer), and RPE1 (immortalized retinal pigment epithelium, Δp53 background) exhibited robust induction in response to MA2 (Figure 5G). However, H2087 (non-small cell lung cancer), HCT116 (colorectal cancer) and MDA-MB-231 (breast cancer) showed limited responsiveness to MA2 (Figure 5G), despite expressing substantial amount of MDA5 (Figure S8F). The cell line-dependent variability of the MA2 responsiveness was also confirmed by IFNβ ELISA (Figure S8G).

Because MA2 binds filamentous MDA5, we hypothesized that the abundance of endogenous dsRNA––and thus pre-existing MDA5 filaments––may determine cellular responsiveness. We therefore asked whether anticancer therapies known to increase endogenous dsRNA and activate RLRs^17-20,25,26,41^ could sensitize otherwise less-responsive cells, such as H2087 cells, to MA2. Pretreatment of H2087 cells with multiple dsRNA-inducing therapies, including DNA polymerase inhibitor aphidicolin (Aph) together with ATR inhibitor (VE-821), MPS1 inhibitors (BAY-1217389, AZ3146), DDX3 inhibitor (RK-33) and METTL3 inhibitor (STM3006), potentiated the MA2-response of H2087 cells, albeit to varying degrees (Figure 5H).

To further establish dsRNA as the key determinant of MA2 responsiveness, we employed A549 cells lacking ADAR1, an A-to-I editing enzyme that acts on dsRNA, because deletion of ADAR1 increases endogenous dsRNA that can activate MDA5^42-44^. Since endogenous MDA5 expression in these cells was insufficient to generate a robust signaling response, we introduced doxycycline (DOX)-inducible MDA5 expression. Consistent with the established role of ADAR1, ADAR1 knockout increased antiviral signaling in a manner dependent on DOX-induced MDA5, as measured by *IFNB* or *CCL5* mRNA (Figure 5I). MA2 further potentiated this response in a manner dependent on DOX-induced MDA5 expression and ADAR1 deficiency (Figure 5I). Further supporting the importance of endogenous dsRNA, the signaling activity of WT MDA5 upon transient overexpression in 293T ADAR1-deficient cells, was dependent on its ability to bind RNA; filament-deficient variant H927A showed no basal signaling and no response to MA2 (Figure 5J). These findings indicate that MA2 does not activate MDA5 *de novo*, but instead potentiates signaling from pre-existing filaments assembled on endogenous dsRNA.

Together, these findings demonstrate that MA2 is a potent context-dependent agonist of MDA5 and identify endogenous dsRNA as a key determinant of cellular responsiveness to MA2, providing a mechanistic rationale for combining MA2 with dsRNA-inducing therapies.

## Discussion

Decades of research have established strong genetic and mechanistic links between MDA5 and a broad range of pathologic and therapeutic processes, including immune disorders, viral infections, mRNA vaccination and cancer therapy. Yet, small-molecule modulators of MDA5 have remained elusive. MDA5 belongs to the SF2 helicase family and possesses an ATP-binding pocket. However, unlike protein kinases, helicase ATP sites are generally shallow, highly polar, and poorly suited for pharmacological intervention. As such, few helicases have proven tractable drug targets, and those that have are typically modulated through allosteric rather than ATP-competitive mechanisms^45-47^. MDA5 presents an additional challenge because its signaling activity depends on cooperative filament assembly on dsRNA, involving extensive protein-protein and protein-RNA interfaces that are difficult to target with conventional small molecules. Here, we exploited this distinctive feature of MDA5 by targeting a cryptic pocket within the filament interface. MA2 stabilizes the signaling-competent filamentous state of MDA5 and enhances antiviral signaling, demonstrating that higher-order protein assemblies can be tractable targets for small molecule-modulation.

Our findings also reveal a unique mode of action for molecular glues, which are increasingly recognized as a powerful mechanism for modulating protein functions^48,49^. Whereas most molecular glues stabilize relatively static protein-protein interfaces, MA2 engages an interface that undergoes substantial conformational remodeling during the ATPase cycle (Figure 6A). ATP binding and hydrolysis normally drive changes in filament geometry that promote filament destabilization and turnover, thereby limiting MDA5 signaling. While MA2 binding attenuates ATP hydrolysis, it does not abolish ATPase activity or ATP-associated conformational changes. Instead, MA2 remains bound to the filament interface while both accommodating and reshaping ATP-associated conformational transitions (Figure 6A). This dynamic interaction enables MA2 to strengthen MDA5-dsRNA and inter-protomer interactions without preventing conformational cycling. Thus, MA2 represents a previously unrecognized class of dynamic molecular glues: small molecules that remain associated with conformationally plastic interfaces while both accommodating and reshaping its structural changes and their functional consequences. Given that many innate immune signaling molecules function through dynamic higher-order assemblies and their conformational cycling^50^, dynamic molecular glues may represent a new strategy for targeting these assemblies.

**Figure 6.**
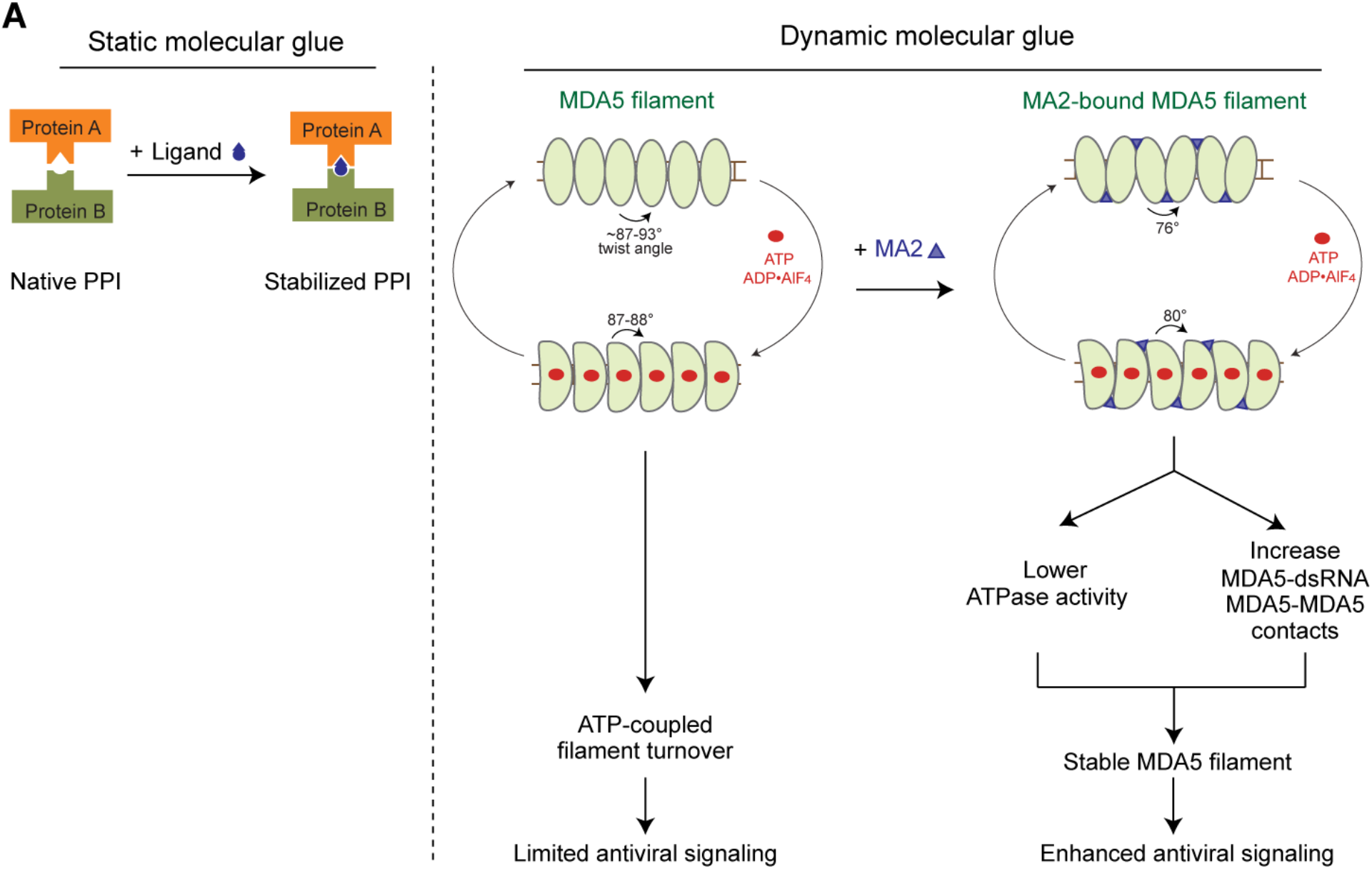
Cartoon model illustrating MA2 as a dynamic molecular glue in contrast to conventional static molecular glues. (A) Static molecular glue: a rigid ligand locks two proteins into a single, fixed protein-protein interface— either by bringing two non-natively interacting proteins into proximity, or by converting a weak, native interaction into a stabilized one — without accommodating further conformational change. Dynamic molecular glue: MA2 engages a structurally plastic interface across multiple nucleotide states of the MDA5 filament. In the Apo state, the filament adopts a high helical twist-dominant conformation; MA2 binding remodels this into a low-twist conformation, increasing MDA5-dsRNA interactions. In the ADP•AlF4-bound state, MA2 accommodates the associated conformational change, and remains bound throughout, attenuating ATP hydrolysis. Both effects converge to stabilize the MDA5 filament, resulting in enhanced antiviral signaling.

Our study also identifies dsRNA as a key determinant of MA2 activity, providing a potential mechanism for tissue and disease selectivity. Because MA2 requires pre-existing dsRNA, its activity is naturally biased toward infected or stressed cells that accumulate exogenous or endogenous dsRNA. This property raises the possibility that MA2 could be used prophylactically to enhance antiviral immunity preferentially upon infection, rather than constitutively inducing systemic inflammation. MA2 may also serve as an adjuvant for mRNA vaccines by selectively enhancing MDA5 activity, which has been shown to contribute to vaccine efficacy^51^, without activating other dsRNA-responsive pathways that suppress antigen expression^52^. In cancer, where genotoxic stress, epigenetic dysregulation, and viral mimicry frequently increase dsRNA abundance, MA2 may preferentially amplify innate immune signaling within tumors, while minimizing systemic toxicity. This selective property also provides a strong rationale for combination strategies with chemotherapeutic or epigenetic agents known to increase intracellular dsRNA accumulation.

Altogether, our study establishes MDA5 as a tractable target for small-molecule modulation and provides a foundation for the future development of both agonists and antagonists. Its dsRNA-dependent activity distinguishes MA2 from other innate immune activators, including STING agonists^6,7^ and PTPN1/2 inhibitors^8^. More broadly, the concept of dynamic molecular glue expands the repertoire of druggable targets to include conformationally flexible protein assemblies and interfaces previously considered undruggable.

## Acknowledgements

We thank Odyssey Therapeutics for sharing the compounds and all members of the Hur lab for their helpful discussions and feedback. This study was supported by Expect Miracles Foundation and the Robert A. and Renee E. Belfer Family Foundation (CP), and Howard Hughes Medical Institute (SH). Cryo-EM data were collected at Cryo-EM facilities at Janelia, Harvard Medical School and University of Massachusetts Medical School. Cryo-EM data processing was supported by the SBGrid and E3 computational platforms.

## Author Contributions

LY determined structures, LY, SP, CT performed experiments, and ZY performed computational analyses. TH generated reagents. ZL, KH and EI assisted NK cell killing experiments. CP, SC, DB, SH supervised the project. LY and SH wrote the manuscript.

## Data availability

The atomic coordinates have been deposited in the Protein Data Bank with accession codes PDB: 37NW, 37NX, 37NY, and 37NZ. The cryo-EM maps included in this study have been deposited in the Electron Microscopy Data Bank under the accession codes EMD-78333, EMD-78334, EMD-78335, EMD-78336. The raw sequencing and read count data were deposited in GSE342766. The scripts used in this manuscript were documented in this Github link: (https://github.com/AnneyYeZiqing/hur_lab_ma2/).

## Supplementary Materials

Figs. S1 to S8; Tables S1 to S3

## Material and Methods

### Double-stranded RNA (dsRNA)

All dsRNAs used in this study were generated by *in vitro* T7 transcription as described previously^30^. DNA templates for RNA synthesis were prepared by PCR amplification. The 512 bp dsRNA sequences were derived from the first 500 bp of the human MDA5 gene, flanked by 5′-gggaga and 5′-tctccc sequences as in the previous study^30^.

### Cell Lines

HEK293T, MCF-7, MDA-MB-231, and RPE1Δp53 cells were maintained in Dulbecco’s Modified Eagle Medium (DMEM) supplemented with 10% fetal bovine serum (FBS). A549, HCC2429, H2087, and H1650 cells were cultured in Roswell Park Memorial Institute (RPMI) 1640 medium supplemented with 10% FBS. HCT116 cells were maintained in McCoy’s 5A (Modified) medium supplemented with 10% FBS. Calu-3 human lung adenocarcinoma epithelial cells (ATCC HTB-55) were cultured in Minimum Essential Medium (MEM; Gibco) supplemented with 10% (v/v) FBS, 1% (v/v) penicillin/streptomycin, 1% (v/v) GlutaMAX, and 1% (v/v) non-essential amino acids. Unless otherwise indicated, all cell lines were maintained under standard culture conditions (37°C in a humidified atmosphere containing 5% CO_2_) and were routinely tested for mycoplasma contamination. MDA5 knockout cell lines were generated by electroporation of a ribonucleoprotein (RNP) complex comprising guide RNA (gRNA) and Alt-R® S.p. Cas9 Nuclease V3 (Integrated DNA Technologies) using the Neon® Transfection System (Thermo Fisher Scientific) and the following parameters: 1,400 V, 20 ms, two pulses for MCF-7 and 1,650 V, 30 ms, one pulse for HCC2429 cell lines. Knockout efficiency was confirmed by Western blotting. For HCC2429 cells, a polyclonal knockout population was used, whereas single-cell knockout clones were isolated for MCF-7 cells. A549-ADAR1 knockout (KO) doxycycline-inducible MDA5 knock-in (A549-ADAR1KO-dox-MDA5) cells were generated by CRISPR/Cas9-mediated knockout of ADAR1, followed by lentiviral transduction with a doxycycline-inducible MDA5 expression construct. 293T-ADAR1KO cells were generated by CRISPR/Cas9-mediated knockout of ADAR1. For NanoBRET assay, A549 cells lacking MAVS and RNase L were used because dsRNA transfection led to significant cell death in a MAVS- and RNase L-dependent manners. Cells were sequentially transduced with the lentivirus harboring doxycycline-inducible MDA5-Halo and stably-expressing NanoLuc-TRIM65 constructs, followed by blasticidin and puromycin selection, respectively. The complete list of gene-specific gRNA sequences is provided in Table S1.

### Plasmids

Mammalian expression plasmids encoding human MDA5 (WT, G495R, H927A, R337G), Plasmids for recombination expression of human MDA5 and human N-terminal CARD truncation variants (ΔCARD) of MDA5 (WT, G495R), RIG-I, LGP-2, mouse Mda5 were generated as previously described^31,36^. Point mutants (C407Y, F373A, R374A, R386A, mMda5-Y408C) were generated by site-directed mutagenesis. MDA5-Halo was cloned into the pInducer20-eGFP-BlasticidinR vector, and NanoLuc-TRIM65 was cloned into the pGK lentiviral vector for stable expression.

### Protein expression and purification

MDA5 full-length protein was expressed in Escherichia coli and purified as previously described^30^ for the Heparin chase assay. Unless otherwise indicated, N-terminal CARD truncation variants (ΔCARD) of MDA5 were used in most other biochemical assays, as ΔCARD retains dsRNA binding, filament formation and MA1/MA2 binding, while exhibiting greater protein stability. The MDA5 were expressed as N-terminally His6-SUMO-tagged proteins in Escherichia coli Rosetta2 (DE3) (Novagen) by induction with 0.25 mM isopropyl β-D-thiogalactopyranoside (IPTG) at 18°C overnight. Cells were lysed by Emulsiflex or sonication and clarified by centrifugation. The proteins were purified using Ni–NTA agarose chromatography, and eluted with Ulp1 protease that cleaves the His_6_-SUMO-tag. The eluted proteins were further purified by chromatography on HiTrap Heparin and Superdex 200 Increase 10/300 columns in buffer containing 20 mM HEPES (pH 7.5), 150 mM NaCl, and 2 mM DTT. mMda5(Δ2CARD) and RIG-I proteins were expressed in Escherichia coli and purified as described before^32^. LGP2 protein was expressed in Sf9 cells using the Bac-to-Bac baculovirus expression systems (Thermo Fisher Scientific) and purified as described before^32^.

### ATPase assay

ATP hydrolysis activity was measured using the Green Reagent (Enzo Life Sciences) according to the manufacturer’s instructions. Purified MDA5Δ2CARD proteins (wild type and indicated mutants) at indicated concentrations were pre-incubated with 512 bp dsRNA (1 ng/µL) and the indicated concentrations of compounds in assay buffer containing 20 mM HEPES (pH 7.5), 150 mM NaCl, and 1.5 mM MgCl_2_, 2mM DTT at room temperature. The reaction was initiated by adding ATP to a final concentration of 2 mM, followed by incubation at 37 °C for the indicated time. Reactions were terminated by the addition of 83.3 mM EDTA. Green Reagent (90 µL) was then added to each reaction mixture at a 9:1 (reagent : reaction) ratio. The release of inorganic phosphate was quantified by measuring absorbance at 650 nm (OD650) using a Synergy2 microplate reader (BioTek). Purified RIG-I (50nM) and LGP (100nM) were analyzed under the same conditions.

### Thermal shift assay

Thermal stability was assessed using a fluorescence-based thermal shift assay. Purified MDA5Δ2CARD proteins (wild type and indicated mutants), mMda5 (WT, Y408C), RIG-I, and LGP2 (0.1 mg/mL) were pre-incubated with MA1/MA2 (200 µM) in the presence or absence of 512 bp dsRNA (50 ng/µL) and ADP•AlF_4_ (0.5 mM) in buffer containing 20 mM HEPES (pH 7.5), 150 mM NaCl, and 1.5 mM MgCl_2_, 2mM DTT. Protein Thermal Shift− Dye (Thermo Fisher Scientific) was added at a 1:1000 dye-to-reaction volume ratio. Thermal melting curves were recorded using a CFX384 real-time PCR detection system (Bio-Rad), and melting temperatures (Tm) were determined from the first derivative of the fluorescence melting curves.

### Heparin chase assay

ATP-driven disassembly of MDA5 filaments was assessed using a heparin chase assay in buffer containing 20 mM HEPES (pH 7.5), 150 mM NaCl, and 1.5 mM MgCl_2_, 2mM DTT. Purified full-length MDA5 proteins (2 µM) were pre-incubated with 512 bp dsRNA (2.4 µg/mL) to form MDA5 filaments under saturating dsRNA conditions in the presence or absence of MA1/MA2 (100 µM) for 10 min at room temperature. The reaction was initiated by the addition of ATP and heparin to final concentrations of 2 mM and 50 µg/mL, respectively, followed by incubation at 37 °C. Reactions were quenched on ice at the indicated time points and analyzed by native gel electrophoresis.

### NanoBRET assay

A549 cells stably expressing TRIM65-NanoLuc and dox-inducible MDA5-Halo were cultured in Opti-MEM with 4% FBS in 96-well plate, as recommended by the manufacturer, and treated with or without MA2 (50 µM) and transfected with or without 512 bp dsRNA (0.5 µg per well). NanoBRET signals were measured at the indicated time points using a Synergy 2 microplate reader (BioTek) equipped with dual-emission filters optimized for NanoBRET detection. Luminescence and fluorescence signals were recorded at 450 nm (bandpass [BP] 80 nm) for the NanoLuciferase donor signal and 610 nm (long-pass [LP]) for the HaloTag acceptor signal, respectively. Corrected NanoBRET ratios were calculated and analyzed according to the manufacturer’s instructions for the NanoBRET− Nano-Glo® Detection Systems (Promega).

### ELISA

Cytokine secretion was assessed by ELISA using cell culture supernatants. For IFNβ analysis, HCC2429 and MCF-7 cells were seeded in 24-well plates at 50% confluency and treated with compounds at the indicated concentrations in medium supplemented with 2% FBS. At 42 h post-treatment, fresh culture supernatants were collected and analyzed using the LumiKine− Xpress hIFN-β 2.0 kit (InvivoGen, luex-hifnbv3) according to the manufacturer’s instructions. For CXCL10 analysis, cells were seeded in 24-well plates at 50% confluency and treated with MA2 in medium supplemented with 10% FBS. At 72 h post-treatment, culture supernatants were collected and analyzed using a human CXCL10 Quantikine ELISA kit (R&D Systems, DIP100) according to the manufacturer’s instructions. For IFNγ analysis, cells were seeded in 24-well plates at 50% confluency in medium supplemented with 10% FBS, and treated with MA2 and/or co-cultured with NK cells as indicated. At 48 h post-treatment, culture supernatants were collected and analyzed using a human IFNγ Quantikine ELISA kit (R&D Systems, DIF50C) according to the manufacturer’s instructions.

For all ELISA assays, absorbance was measured using a Synergy M1 microplate reader (BioTek), and cytokine concentrations were calculated based on standard curves generated from recombinant cytokine standards.

### Immunoblotting

Cells were lysed in SDS lysis buffer containing 1% SDS, 50 mM Tris-HCl (pH 7.5), 150 mM NaCl, 10 mM DTT, and 5 mM EDTA, followed by boiling at 95 °C for 10 min. Protein concentrations of clarified lysates were determined using a BCA protein assay kit (Thermo Fisher Scientific). Equal amounts of total protein were mixed with 1× SDS sample buffer containing 62.5 mM Tris-HCl (pH 6.8), 10% glycerol, 2% SDS, and 2.5% β-mercaptoethanol, and resolved on 4–15% Tris-glycine gels (Bio-Rad, 4561086). Proteins were transferred onto 0.45 µm PVDF membranes (Thermo Fisher Scientific, 88518). Membranes were blocked with 5% (w/v) nonfat dry milk in PBST (PBS containing 0.1% Tween-20) for 30 min at room temperature. Blots were incubated with primary antibodies diluted in PBST supplemented with 1% BSA overnight at 4 °C with gentle agitation, followed by incubation with HRP-conjugated secondary antibodies diluted in PBST supplemented with 3% milk for 1 h at room temperature. Signals were detected using a ChemiDoc MP Imaging System (Bio-Rad). The following primary antibodies were used: MDA5 (Cell Signaling Technology, D74E4), phospho-STAT1 (Cell Signaling Technology, 9167), β-actin (Cell Signaling Technology, 8457), Flag (Sigma/Millipore, F4049), ADAR1 (Santa Cruz, sc-73408), SARS-CoV-2 Spike (IgG1 CR3022), IAV (IgG1 70-1F02). Immunoblotting experiments performed using the Jess Automated Western System (ProteinSimple Technology, Part number 004-650) according to the manufacturer’s instructions were specified in the corresponding figure legends.

### RT-qPCR

Total RNA was extracted using the Direct-zol RNA Miniprep Kit (Zymo Research, c1078-250), followed by DNase I digestion for 30 min at room temperature. cDNA was synthesized using the High-Capacity cDNA Reverse Transcription Kit (Life Tech) according to the manufacturer’s instructions. Real-time PCR was performed using gene-specific primers or random primers, SYBR Green Master Mix (Applied Biosystems), and a Real-Time PCR System (Bio-Rad). The complete list of gene-specific primers is provided in Table S2.

### RNA-seq

MCF-7 and HCC2429 cells were cultured in 24-well plates in DMEM or RPMI, respectively, supplemented with 10% FBS, and treated with DMSO, 50 µM MA2, or 10 µM diABZI, as indicated. To achieve comparable MA2- and poly(I:C)-induced IFNβ signaling, MCF-7 cells were cultured in DMEM supplemented with 2% FBS to enhance MA2 efficiency under low-serum conditions, and 10 ng/mL poly(I:C) were transfected. Cells were harvested 24 h after treatment by direct lysis in TRIzol reagent, and total RNA was isolated using the Direct-zol RNA Miniprep Kit (Zymo Research), including on-column DNase I digestion. Purified RNA was used for cDNA library preparation with the SMARTer Stranded Total RNA-Seq Kit v3 Pico Input Mammalian (Takara Bio) according to the manufacturer’s instructions. Library quality was assessed using an Agilent TapeStation, and libraries were sequenced on an Illumina NovaSeq 6000 platform with 150-bp paired-end reads (2 × 150 bp), generating approximately 15 million paired-end reads per sample.

### Bioinformatics analysis

Paired-end sequencing reads were trimmed using Trimmomatic and aligned to the human reference genome (GRCh38.p13) using STAR. Coordinate-sorted BAM files were generated with default alignment and multimapping parameters. Gene-level read counts were generated using featureCounts (Subread) against GENCODE v43 exon annotations in paired-end mode. Multi-mapping reads were not counted. Differential gene expression analysis was subsequently performed in R using DESeq2. DESeq2-generated normalized count matrices and differential expression statistics were used for downstream expression analyses and visualization. Adjusted p-values for pathway enrichment are calculated by EnrichR.

### IFNβ reporter dual luciferase assay

ADAR1-deficient HEK293T cells were cultured in 48-well plates in DMEM supplemented with 10% FBS. At approximately 80% confluency, cells were co-transfected with a pFLAG-CMV4 plasmid encoding MDA5 (10 ng), an IFNβ promoter-driven firefly luciferase reporter plasmid (100 ng), and a Renilla luciferase control plasmid (pRL-CMV, 5 ng) using Lipofectamine 2000 (Life Technologies) according to the manufacturer’s instructions. After 4 h of transfection, the medium was replaced with fresh DMEM containing 2% FBS, supplemented with or without MA2 (50 µM). Cells were lysed 20 h after treatment, and IFNβ promoter activity was measured using the Dual-Luciferase Reporter Assay System (Promega) with a Synergy 2 microplate reader (BioTek). Firefly luciferase activity was normalized to Renilla luciferase activity.

### Viral infection assays

Encephalomyocarditis virus (EMCV; ATCC VR-129B) was obtained from ATCC. For EMCV infection, MCF-7 or HCC2429 cells were pre-treated with DMSO or MA2 (50 µM) for 6 h in medium supplemented with 2% FBS, and then infected with EMCV at the indicated MOIs. Virus adsorption was carried out for 1 h. Following adsorption, fresh medium was added to restore the final FBS concentration to 10%, and cells were cultured for an additional 24 h before harvesting for RT-qPCR analysis. For SARS-CoV-2 and influenza A virus (IAV) infection, Calu-3 cells were plated at a density of 45,000/well in black 96 well collagen coated plates (Corning), and were pre-treated with vehicle or MA2 (50 µM) for 6 h and infected with the SARS-CoV-2 WA.1 strain (BEI Resources, NR-52281; USA-WA1/2020) or influenza A virus (A/Puerto Rico/8/1934 [PR8] strain; courtesy of Dr. Scott Hensley) at the indicated MOIs for 48 h. For automated microscopy, infected cells were fixed with 4% formaldehyde for 15 min at room temperature and washed three times with PBS. Cells were permeabilized and blocked with 2% BSA in PBS containing 0.1% Triton X-100 (PBS-T) for 1 h at room temperature. Plates were incubated overnight at 4 °C with either anti-SARS-CoV-2 Spike antibody (CR3022, IgG1) or anti-IAV antibody (70-1F02, IgG1), washed three times with PBS-T, and incubated with Alexa Fluor-conjugated secondary antibodies and Hoechst 33342 for 1 h at room temperature. After three additional washes with PBS, plates were sealed and imaged using an ImageXpress Micro automated microscope (Molecular Devices). Images were acquired with a 10× objective from nine fields per well. Total and infected cell numbers were quantified using MetaXpress software (v5.3.3), and the percentage of infected cells was calculated.

### NK cell killing assay

Human NK Cell Isolation Kit (STEMCELL) from anonymized healthy donor–derived peripheral blood mononuclear cells, which were obtained through Crimson Core at Brigham and Women’s Hospital under Dana-Farber Cancer Institute protocol NHSR#428198. Isolated NK cells were expanded for 14 days in NK MACS Medium (Miltenyi Biotec) supplemented with 5% human serum (Sigma-Aldrich) and human IL-2 (250 ng/mL, R&D Systems) prior to use. The medium was refreshed every 3 days. Cell viability was assessed using the CellTiter-Glo® Luminescent Cell Viability Assay (Promega, Cat. No. G7571), and cytotoxicity was determined based on the reduction in cell viability, according to the manufacturer’s instructions.

For the NK-tumor co-culture experiments, cancer cells were seeded in 96-well tissue culture plates at a density of 3000 (MCF-7) or 10,000 (HCC2429) cells per well. After treatment with MA2 and/or co-culture with NK cells (NK-to-target ratio 1:5) for 72 hours, luminescence was measured using a Synergy2 microplate reader (BioTek) to quantify cell viability. IL-2 (250 ng/mL) was maintained in the culture medium throughout the experiment to support NK cell viability and function.

### Cryo-EM sample preparation and data collection

MDA5Δ2CARD-G495R (90 µM) was incubated with 512 bp dsRNA (50 ng/µL) in buffer containing 20 mM HEPES (pH 7.5), 150 mM NaCl, and 1.5 mM MgCl_2_, 2mM DTT at room temperature for 2∼3 hours in the presence or absence of 100 µM MA1 or MA2 and with or without 2 mM ADP·AlF_4_ as indicated. The resulting complexes were cross-linked with 0.5% glutaraldehyde for 10 minutes at room temperature. Samples were applied to freshly glow-discharged C-flat 300-mesh copper grids (CF-1.2/1.3, Electron Microscopy Sciences) and blotted for 5 seconds before plunge-freezing in liquid ethane using a Vitrobot Mark IV (FEI) set at 100% humidity. Grid screening was performed at the Harvard Cryo-EM Center and the UMass Cryo-EM Core Facility using a Talos Arctica (FEI) microscope to identify regions with optimal particle distribution and ice thickness. High-resolution data were collected at the Janelia Cryo-EM Facility using a Titan Krios transmission electron microscope (Thermo Fisher Scientific) operated at 300 kV and equipped with a Gatan K3 direct electron detector. Movies were recorded at a nominal magnification of ×105,000, corresponding to a calibrated pixel size of 0.827 Å. Each movie consisted of 50 frames collected with a total electron exposure of 60 e^−^/Å^2^. Data were acquired over a defocus range of −0.8 to −1.8 µm. A total of 12,506 movies were collected for apo-MDA5 filament, 11,289 movies for MA1-bound filament, 4,680 movies for the MA2-bound filament, and 10,932 movies for the MA2-ADP·AlF_4_-bound filament.

### Cryo-EM data processing and structure refinement

Cryo-EM data were processed using cryoSPARC (v4.2.0) and RELION (v4.0.1). Dose-fractionated movies were motion-corrected using MotionCor2, and contrast transfer function (CTF) parameters were estimated with CTFFIND4 (v4.1).

For the MA1-bound filament dataset, filament coordinates were manually picked in RELION and used to train a Topaz particle-picking model. The trained Topaz model was subsequently applied for automated particle picking. The helical segments were extracted using a box size of 240 pixels with an inter-box distance of 46 Å. A total of 5,356,653 particles were initially extracted and subjected to multiple rounds of reference-free 2D classification, yielding 5,081,531 high-quality particles for ab initio reconstruction. The particles were separated into six ab initio classes, and 1,332,057 particles from the best-resolved class were selected for 3D classification and non-uniform refinement, resulting in a reconstruction with an overall resolution of 3.60 Å with 610,336 particles. The apo-MDA5 filament dataset was analyzed using a box size of 480 pixels and processed using the same workflow, including 2D classification, ab initio reconstruction, 3D classification, and non-uniform refinement.

For the MA2-bound filament dataset, particles were picked using the filament picker implemented in cryoSPARC. 2,882,609 particles were initially extracted (Filament Diameter: 50 Å; Separation distance between segments: 45 Å; Extraction box size: 280 pixels) and subjected to multiple rounds of reference-free 2D classification, yielding 1,440,845 high-quality particles for ab initio reconstruction. The particles were separated into six ab initio classes, and 348,721 particles from the best-resolved class were selected for non-uniform refinement, resulting in a reconstruction with an overall resolution of 2.81 Å. The MA2-ADP·AlF_4_-bound filament and MA2-bound filament datasets were processed using the same workflow of 2D classification, ab initio reconstruction, 3D classification, and non-uniform refinement.

Atomic models were generated by rigid-body fitting the previously reported MDA5–dsRNA crystal structure (PDB: 4GL2) into the cryo-EM density map using UCSF Chimera. Manual model rebuilding was performed in Coot, followed by real-space refinement using phenix.real_space_refine. The MA1 and MA2 ligands were manually built and fitted into the corresponding densities in Coot. For the MA2–ADP·AlF_4_ map, the previously reported MDA5–dsRNA cryo-EM structure (PDB: 7JL2) was rigid-body fitted into the density map and subsequently refined. Model quality was assessed using MolProbity within the PHENIX software suite, and model-to-map agreement was evaluated by calculating model-versus-map Fourier shell correlation (FSC) curves in cryoSPARC. Statistics for cryo-EM data collection, image processing, and model refinement are provided in the Table S3. Structural figures were prepared using PyMOL (Schrödinger) and UCSF Chimera. All cryo-EM image-processing and model-building software were installed and maintained through the SBGrid software collection.

### Protein-RNA interaction analysis

Effect of MA2 on MDA5-RNA interaction was analyzed by a custom Python script (see GitHub), which identified protein-RNA contacts (defined as heavy-atom pairs within 4.5 Å) and calculated, for each protein residue, the total contact count summed across all RNA residues. Differences in total contact count (Δcontact count) were calculated by comparing +MA2 and - MA2 structures. Positive values indicate a net gain of RNA contact upon MA2 binding, while negative values indicate a net loss for that residue. Summing Δcontact counts across all protein residues yielded net increases of +119, +124, +177, +51 for the four protomers (clockwise from the upper left panel in Figure 4D), indicating an overall increase in protein-RNA interactions, particularly in the first three protomers. Similar results were obtained using a more stringent 4.0 Å contact cutoff (+48, +38, +39, -17), with three of the four protomers still exhibiting substantial gains in RNA contacts.

**Figure S1.**
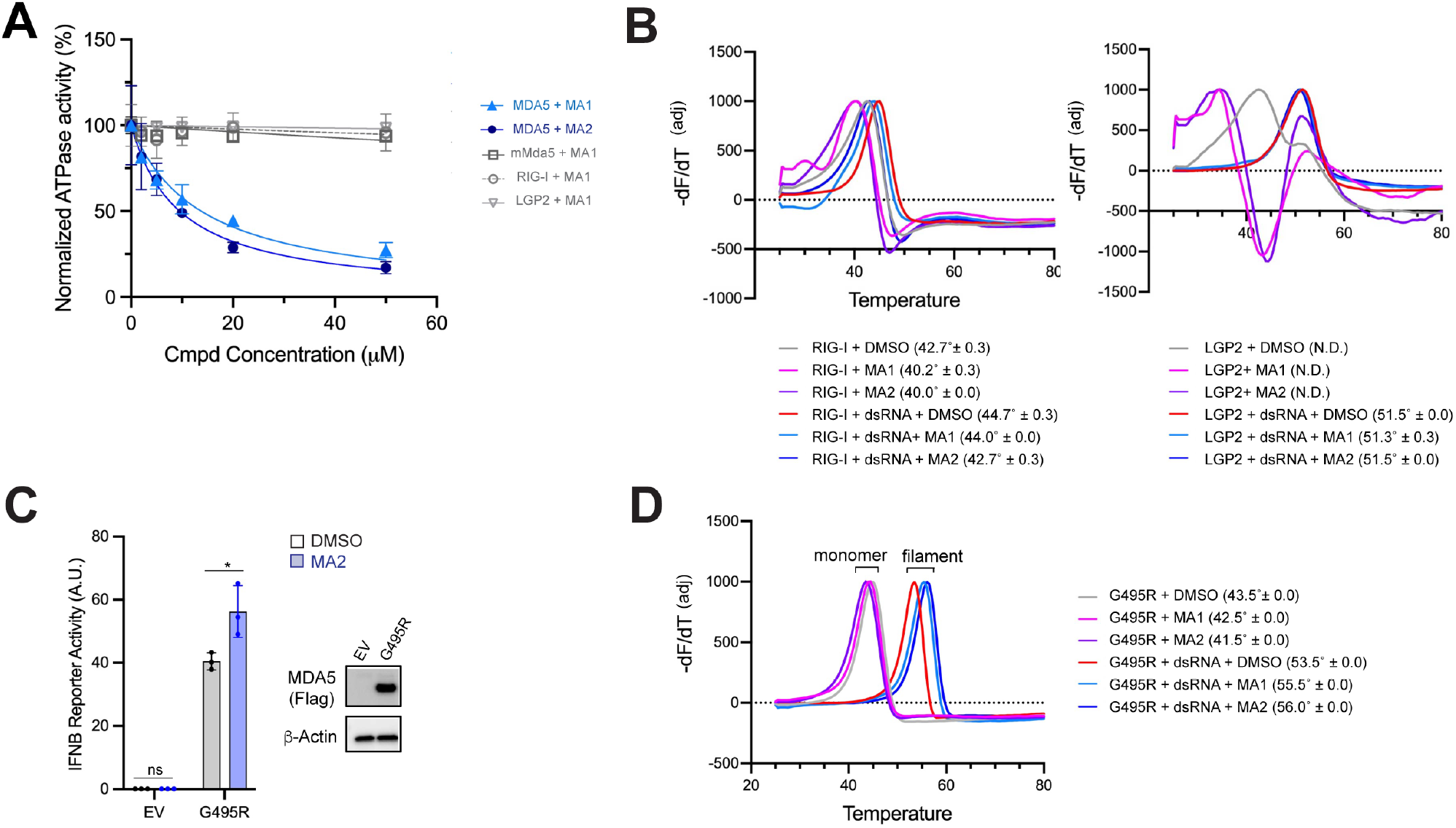
MA1 and MA2 stabilize filamentous MDA5, related to Figure 1. (A) Schematic overview of the antibody cloning process of Effect of MA1/MA2 on the ATPase activity of MDA5. MDA5 (200 nM) was incubated with 512 bp dsRNA (1 ng/μL) and MA1/MA2 (indicated concentration, DMSO-normalized), and reactions were initiated by ATP addition and quenched at 0.5 min. Equivalent condition was used for human RIG-I, LGP2 and mouse Mda5. N-terminal CARD deletion construct was used for human MDA5 and mMda5 as they bind dsRNA and forms filaments similarly to full-length proteins, while displaying greater biochemical stability. Data are shown as average ± S.D. (n=3). (B) Thermal shift assays measuring compound binding to RIG-I and LGP2. RIG-I or LGP2 protein was incubated with MA1 or MA2 (200 μM) in the presence and absence of 512 bp dsRNA (with 5’ppp). The melting temperature (Tm in parenthesis) was calculated from the inflection point of the derivative of fluorescence with respect to temperature (−dF/dT). Amplitude of the derivative was adjusted for visual clarity. Tm was derived from 3 biological repeats (mean±S.D.). (C) IFNβ reporter dual luciferase activity as measured by dual luciferase assay upon transient expression of MDA5 G495R in 293T-ADAR1KO cells in the presence or absence of MA2 (50 μM). EV, empty vector control. Data are shown as average ± S.D. (n=3). \**p* <0.05 was calculated by two-tailed student’s t test. (D) Thermal shift assays measuring compound binding to G495R MDA5. Experiments and analyses were done as in Figure 1E and melting temperature (Tm) is presented as average ± S.D. from 3 biological repeats.

**Figure S2.**
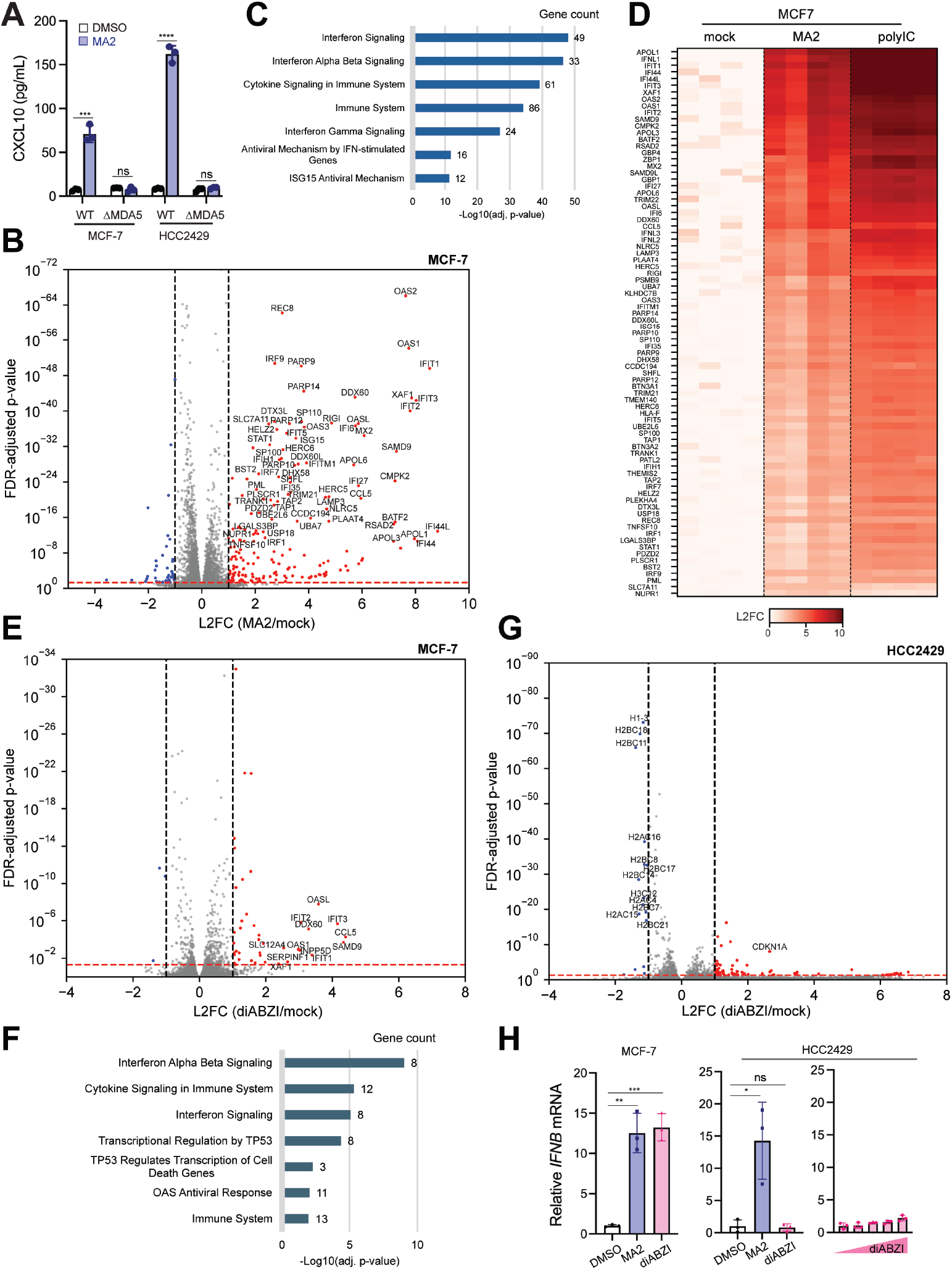
MA2 potentiate MDA5 signaling, related to Figure 2. (A) ELISA of CXCL10 induction in response to MA2 (50 μM) in WT and ΔMDA5 cells. Data are shown as average ± S.D. (n=3). \*\*\*\**p* <0.0001, \*\*\**p* <0.001 were calculated by two-tailed student’s t test. (B) Volcano plot of differentially expressed genes in MCF7 cells upon MA2 stimulation. Significantly upregulated genes (log2-fold change (L2FC) > 1, false discovery rate-adjusted *p*-value (padj) < 0.05) are colored red; significantly downregulated genes (L2FC < -1, padj < 0.05) in blue, and the rest in grey. (C) Reactome pathway enrichment analysis of MA2-upregulated genes (padj < 0.05, L2FC > 1) in MCF-7 cells. The top 7 most significantly enriched categories are shown. Adjusted p-values for pathway enrichment are calculated by EnrichR. (D) Heatmap of gene expression changes for MA2-upregulated genes (padj < 0.05, L2FC > 1 for MCF-7. Cells were stimulated with DMSO (mock), MA2, or polyIC under 2% FBS supplementary. PolyIC (10 ng/mL) was introduced by lipofectamine transfection. (E) Volcano plot of differentially expressed genes in MCF7 cells upon diABZI stimulation, using the same color code as in (B). (F) Reactome pathway enrichment analysis of diABZI-upregulated genes (padj < 0.05, L2FC > 1) in MCF-7 cells. (G) Volcano plot of differentially expressed genes in HCC2429 cells upon diABZI stimulation, using the same color code as in (B). (H) RT-qPCR analysis of *IFNB* mRNA induction in response to MA2 (50 μM) or diABZI (10 μM) in MCF-7 and HCC2429 cells. diABZI beyond 10 μM induced toxicity and does not significantly increase antiviral signaling in HCC2429. Data are shown as average ± S.D. (n=3). \*\*\**p* <0.001, \*\**p* <0.01, \**p* <0.05 were calculated by two-tailed student’s t test.

**Figure S3.**
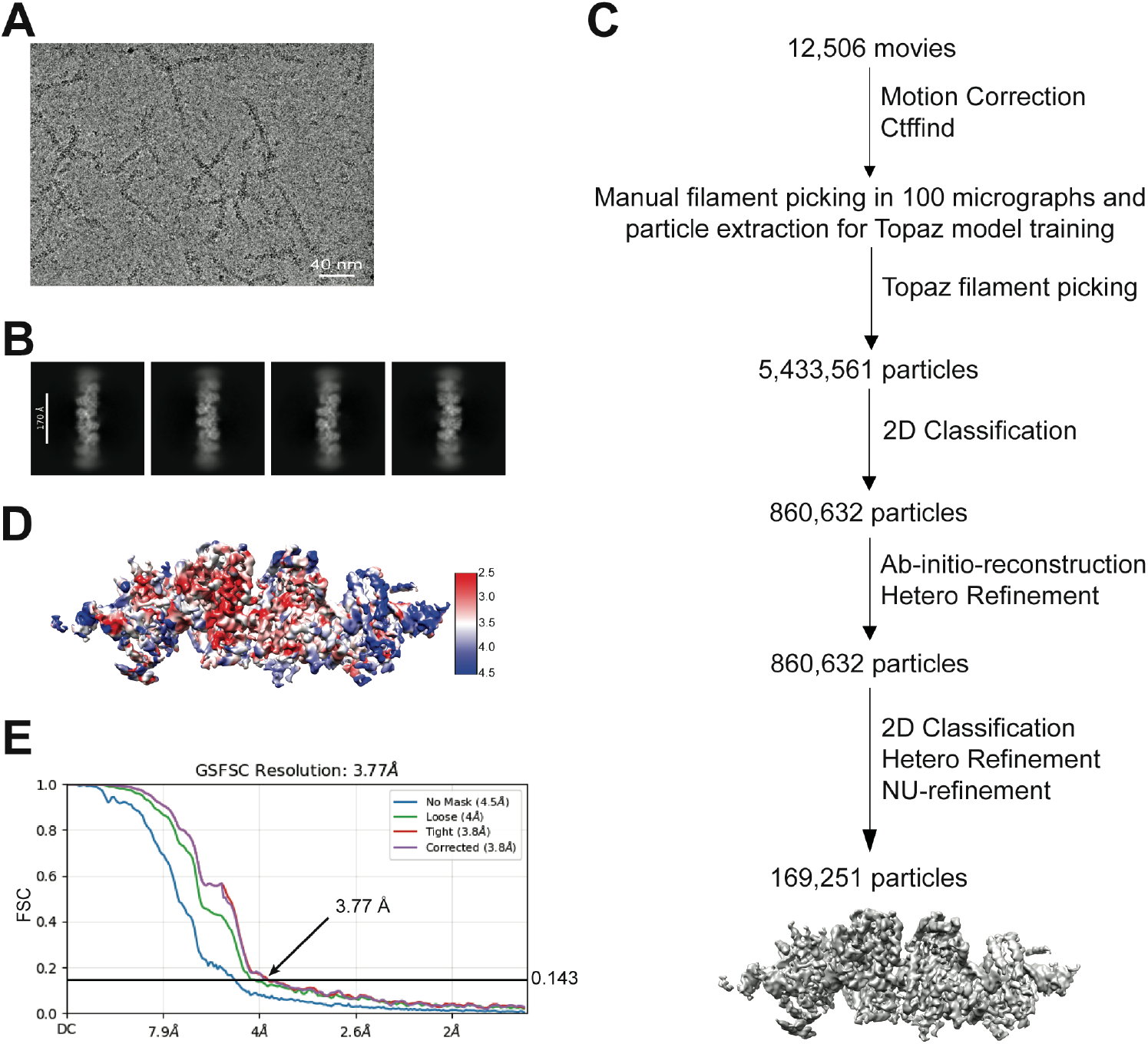
Cryo-EM reconstruction of Apo MDA5 filament, related to Figure 3. (A) Representative cryo-EM micrograph of MDA5 filaments. (B) Representative 2D class averages used for 3D reconstruction of MDA5 filaments. (C) Cryo-EM data processing workflow for the MDA5 filament dataset (see Methods for details). (D) Local resolution map of MDA5 filaments calculated using CryoSPARC. The resolution range is indicated by the color bar. (E) Fourier shell correlation (FSC) analysis of the MDA5 filament reconstruction in CryoSPARC. The overall resolution was determined using the gold-standard FSC = 0.143 criterion.

**Figure S4.**
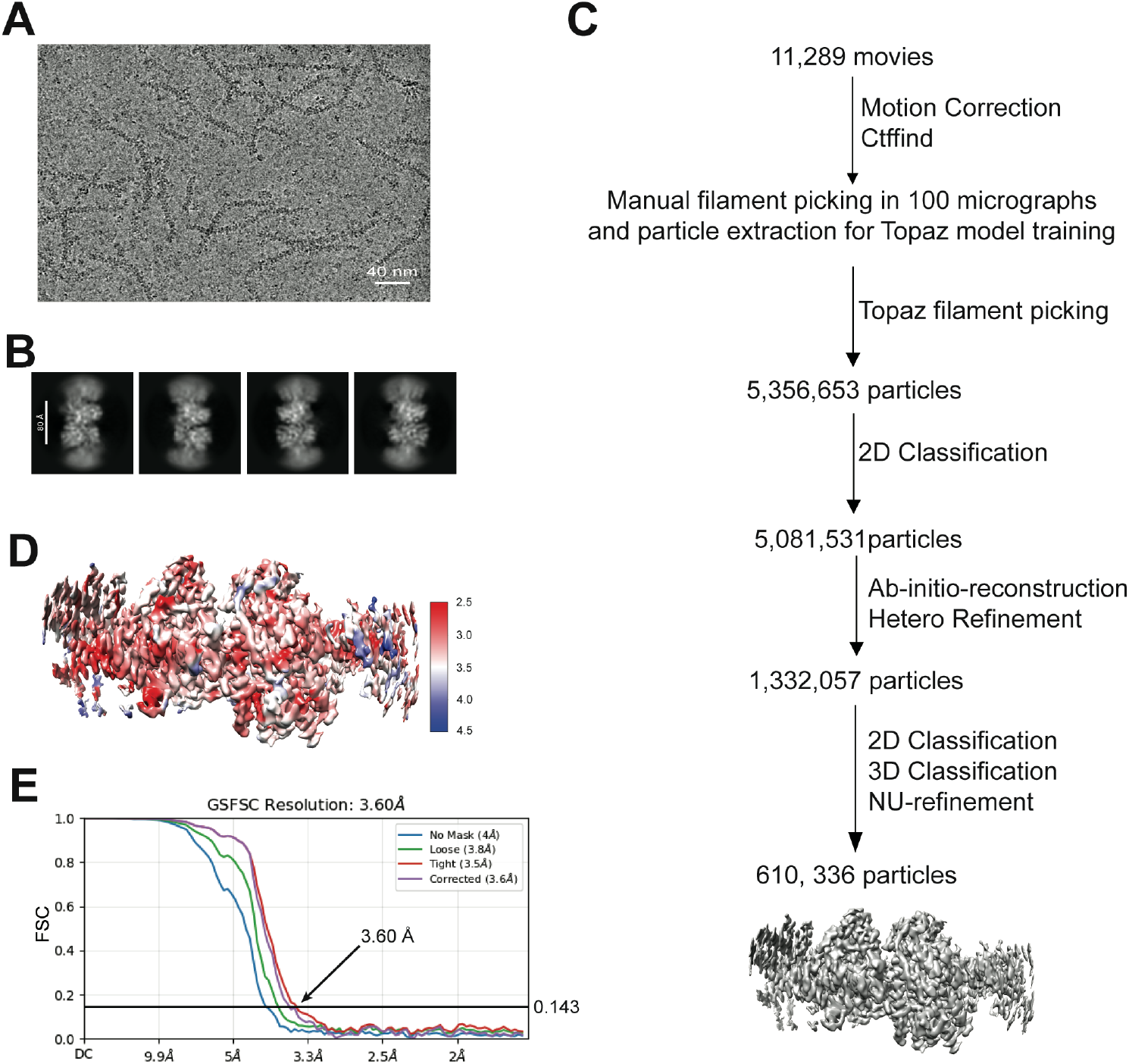
Cryo-EM reconstruction of MA1-bound MDA5 filament, related to Figure 3. (A) Representative cryo-EM micrograph of MDA5 filaments bound to MA1. (B) Representative 2D class averages used for 3D reconstruction of MA1-bound complex. (C) Cryo-EM data processing workflow for the MA1-bound complex dataset (see Methods for details). (D) Local resolution map of MA1-bound complex calculated using CryoSPARC. The resolution range is indicated by the color bar. (E) Fourier shell correlation (FSC) analysis of MA1-bound complex reconstruction in CryoSPARC. The overall resolution was determined using the gold-standard FSC = 0.143 criterion. =====

**Figure S5.**
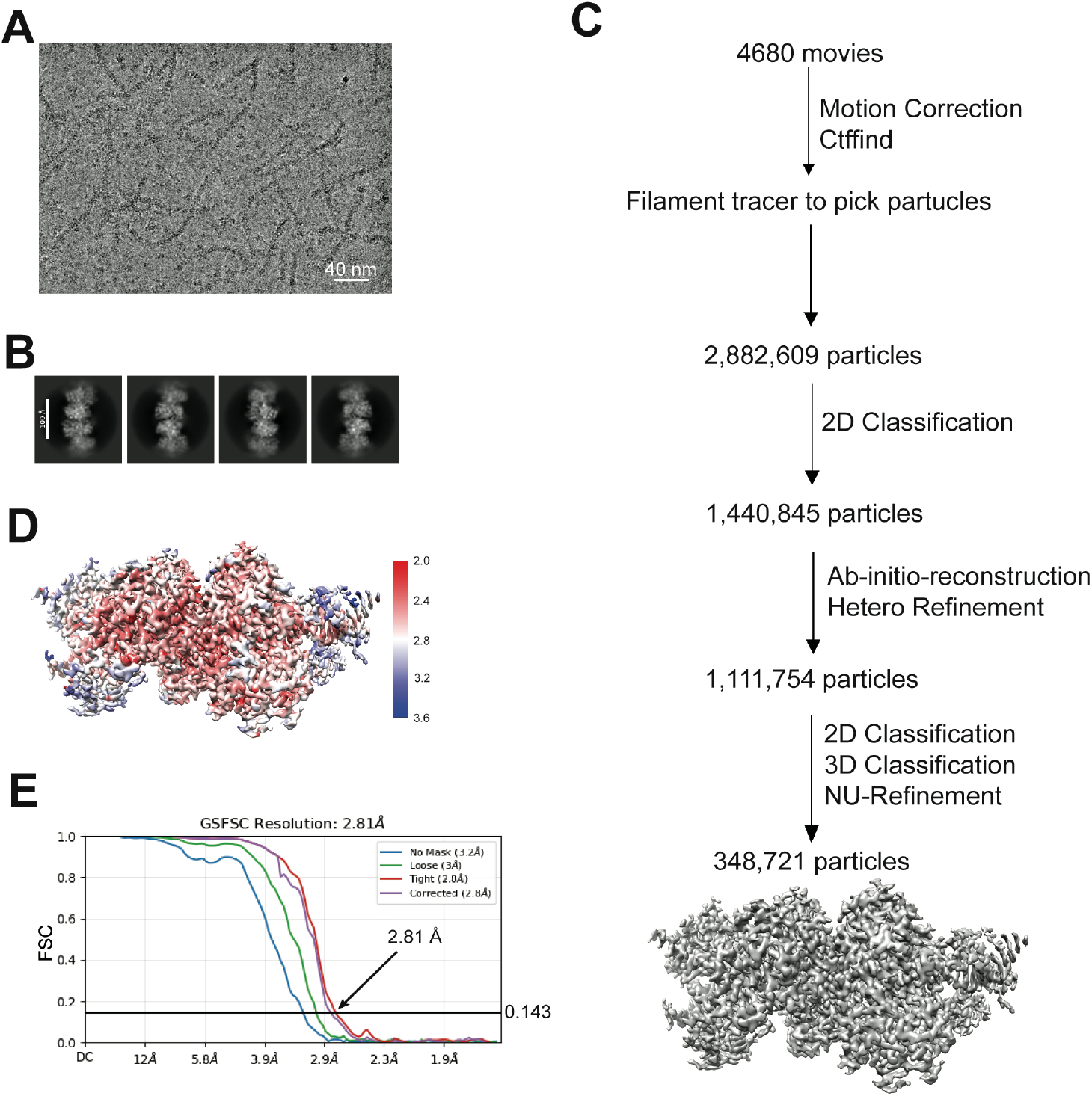
Cryo-EM reconstruction of MA2-bound MDA5 filament without ADP•AlF4, related to Figure 3. (A) Representative cryo-EM micrograph of MDA5 filaments bound to MA2. (B) Representative 2D class averages used for 3D reconstruction of the MA2-bound complex. (C) Cryo-EM data processing workflow for the MA2-bound complex dataset (see Methods for details). (D) Local resolution map of the MA2-bound complex calculated using CryoSPARC. The resolution range is indicated by the color bar. (E) Fourier shell correlation (FSC) analysis of the MA2-bound complex reconstructed in CryoSPARC. The overall resolution was determined using the gold-standard FSC = 0.143 criterion.

**Figure S6.**
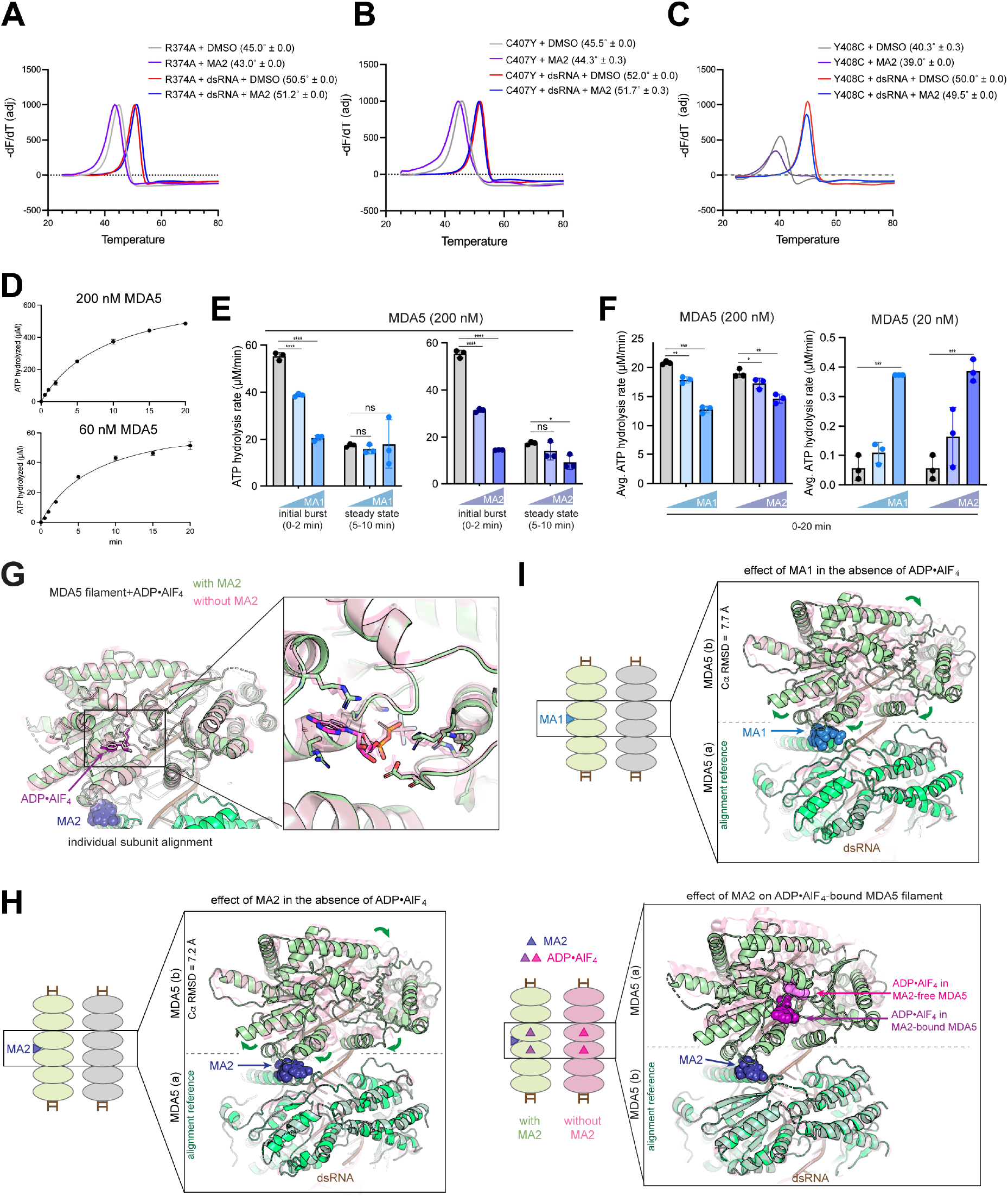
Effect of MA1 and MA2 on MDA5 ATP hydrolysis and filament geometry, related to Figure 4. (A-C) Thermal shift assays measuring compound binding to (A) R374A, (B) C407Y human MDA5 and (C) Y408C mouse Mda5. Indicated protein was incubated with MA1 or MA2 (200 μM) in the presence and absence of 512 bp dsRNA. The melting temperature (Tm in parenthesis) was calculated from the inflection point of the derivative of fluorescence with respect to temperature (−dF/dT). Amplitude of the derivative was adjusted for visual clarity. Tm was derived from 3 biological repeats (mean±S.D.). (D)ATP hydrolysis time course. MDA5ΔCARD (200 nM or 60 nM) was incubated with 512 bp dsRNA (1ng/μL) and the reactions were initiated by ATP addition and quenched at indicated time points. MDA5 hydrolyzes ATP only upon dsRNA binding while ATP hydrolysis in turn destabilizes MDA5 filament. As a result, MDA5 shows high initial burst of ATP hydrolysis (t = 0-2 min) by preassembled filaments, reflecting the intrinsic ATPase activity of the filamentous MDA5. By contrast, the steady-state kinetics (t>5 min) is slower because the apparent ATPase rate reflects both the intrinsic ATPase activity and ATP-driven filament turnover. Data are represented in mean±S.D. from 3 biological repeats. (E)Effect of MA1/2 on the ATPase activity of MDA5. While MA1/2 attenuated the initial burst kinetics, it had a much smaller effect in the steady state due to its filament stabilizing effect, which increases the pool of MDA5 molecules competent for ATP hydrolysis, indirectly counteracting its suppressive effect on the intrinsic ATPase activity. The initial burst rate was determined by linear fit of t=0-2min data points; the steady-state rate was derived from t=5 and 10 time points. \*\*\*\**p* <0.0001, \**p* <0.05 were calculated by two-tailed student’s t test. (F) Effect of MA1/2 on MDA5 ATPase activity under filament-limiting (20 nM MDA5) vs. filament-stable (200 nM MDA5) conditions. MDA5, 512 bp dsRNA (1 ng/μL), and MA1/2 were mixed, reactions were initiated with ATP, and products were quenched and quantified at 20 min. Under filament-limiting conditions, the apparent ATPase activity is influenced more strongly by the filament stabilizing effect of MA1/2 than by their suppression of the intrinsic ATPase activity. Consequently, MA1/2 increased apparent ATP hydrolysis activity under filament-limiting condition, whereas they reduced ATPase activity under filament-stable conditions. \*\*\**p* <0.001, \*\**p* <0.01, \**p* <0.05 were calculated by two-tailed student’s t test. (G) Structural superposition of ADP•AlF4-bound MDA5 filament with (green) and without (pink) MA2, aligned on individual protomers. Inset: close-up view of the ADP•AlF4-binding pocket and surrounding residues, shown as sticks. (H) Effect of MA2 binding to MDA5 filament in the absence (left) or presence (right) of ADP•AlF4 (purple spheres). In each nucleotide state, MA2-bound filament (green) was compared with MA2-free filament (transparent pink). Dimeric segment of MDA5 filament was aligned on protomer (a). Ca RMSD is for protomer (b). The position of ADP•AlF4 in protomer (b) is displaced by ∼11Å between the MA2-free and MA2-bound filaments. (I) Structural alignment of the nucleotide-free MDA5 filament with and without MA1. MDA5 protomer (b) was compared, while aligning on the protomer (a).

**Figure S7.**
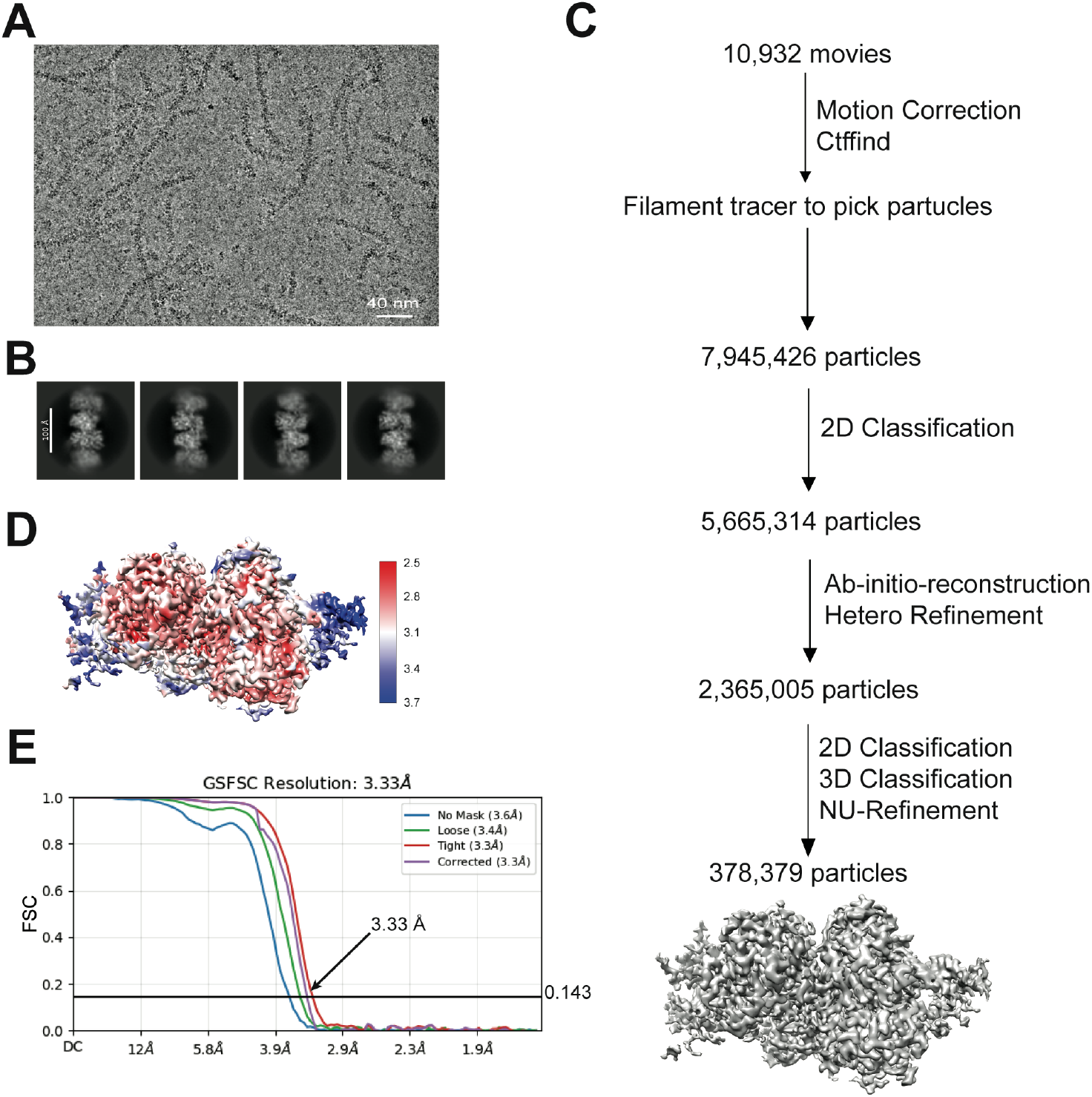
Cryo-EM reconstruction of MDA5 filament bound with MA2 and ADP•AlF4, related to Figure 4. (A) Representative cryo-EM micrograph of MDA5 filaments bound to MA2 in the presence of ADP·AlF_4_. (B) Representative 2D class averages used for 3D reconstruction of the MA2–ADP·AlF_4_-bound complex. (C) Cryo-EM data processing workflow for the MA2–ADP·AlF_4_-bound complex dataset (see Methods for details). (D) Local resolution map of the MA2–ADP·AlF_4_-bound complex calculated using CryoSPARC. The resolution range is indicated by the color bar. (E) Fourier shell correlation (FSC) analysis of the MA2–ADP·AlF_4_-bound complex reconstructed in CryoSPARC. The overall resolution was determined using the gold-standard FSC = 0.143 criterion.

**Figure S8.**
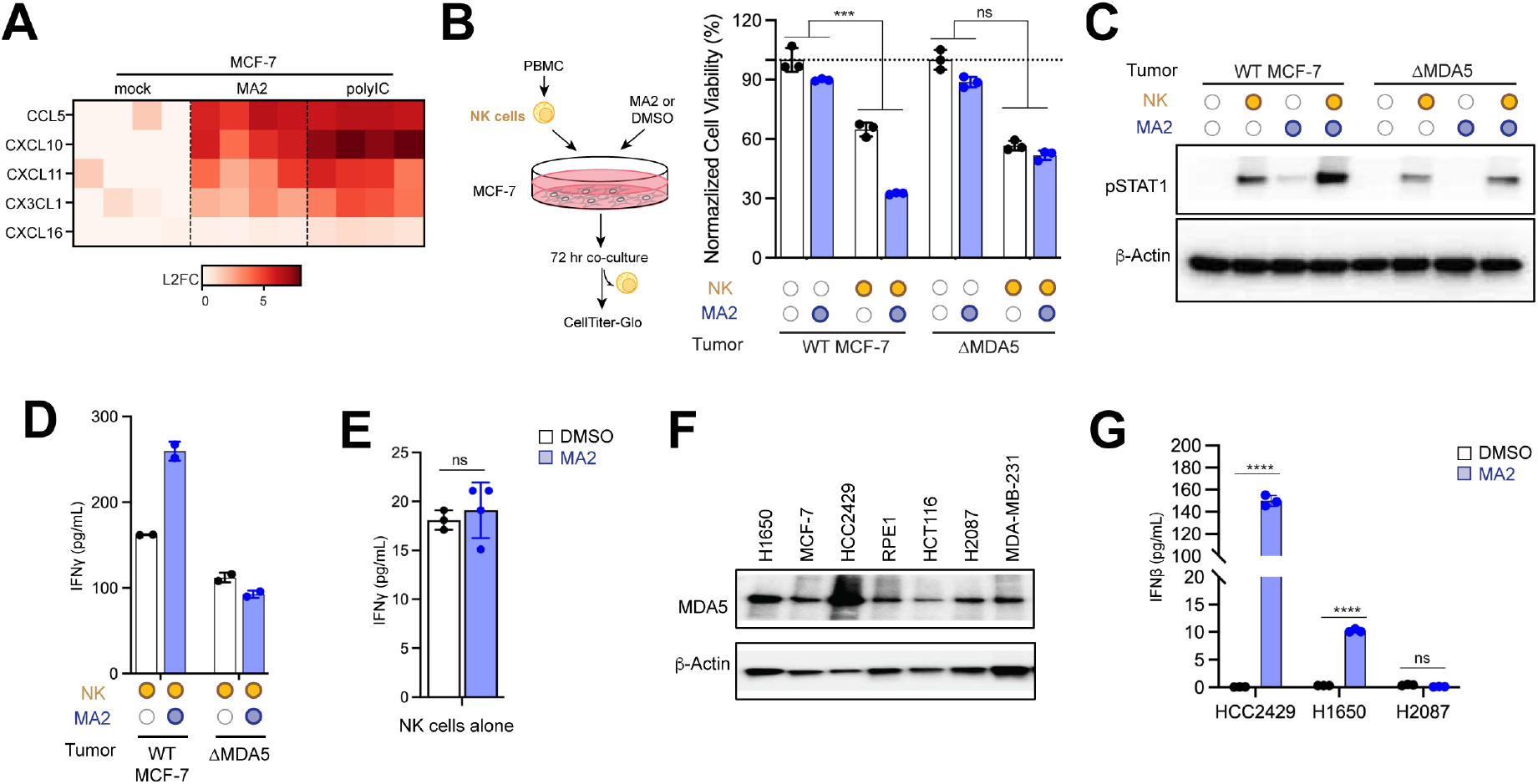
MA2 synergizes with NK cells in antitumor immunity, related to Figure 5. (A) Heatmap of MA2-upregulated CCL and CXCL chemokines (padj < 0.05) for MCF-7 identified in Figure S2D. Cells were stimulated with DMSO (mock), MA2 (50 μM), or polyIC under 2% FBS supplementary. PolyIC (10 ng/mL) was introduced by lipofectamine transfection. (B) Effect of MA2 on NK cell-mediated tumor killing. MCF-7 cells (target) were treated with MA2 (50 μM) and co-cultured with primary human NK cells at an NK-to-target ratio of 1:5 for 72 hrs. MCF-7 cell viability was measured by CellTiter-Glo luminescence, with the NK cell signal subtracted and values normalized to DMSO-treated controls. \*\*\**p* <0.001 was calculated by Two-Way ANOVA analysis, comparing the effect of MA2 on tumor cell killing in the presence versus absence of NK cells, assessing the interaction between MA2 and NK cells. (C) Effect of MA2 on STAT1 phosphorylation in tumor-NK co-culture. MCF-7 cells were co-cultured with NK cells at an NK-to-target ratio of 1:10 in the presence and absence of MA2 (50 μM). After 48 hrs of co-culture, NK cells were removed by PBS washing, and MCF-7 cell lysates were collected for immunoblot analysis. (D) Effect of MA2 on IFNγ expression in tumor-NK co-culture. MCF-7 cells were co-cultured with NK cells for 48 hrs as in (b) and secreted IFNγ was measured by ELISA. (E) Effect of MA2 on IFNγ expression in NK cells. NK cells were incubated with MA2 for 72 hrs prior to IFNγ ELISA. (F) Immunoblot analysis of endogenous MDA5 in indicated cell lines. (G) Effect of MA2 on IFNβ expression in indicated cell lines. Cells were incubated with MA2 for 24 hrs prior to IFNβ ELISA. \*\*\*\**p* <0.0001 was calculated by two-tailed student’s t test.

**Table S1.**
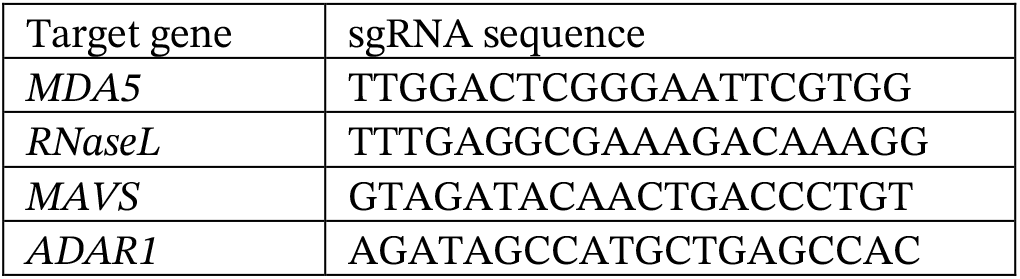
sgRNA sequences for CRISPR-Cas9.

**Table S2.**
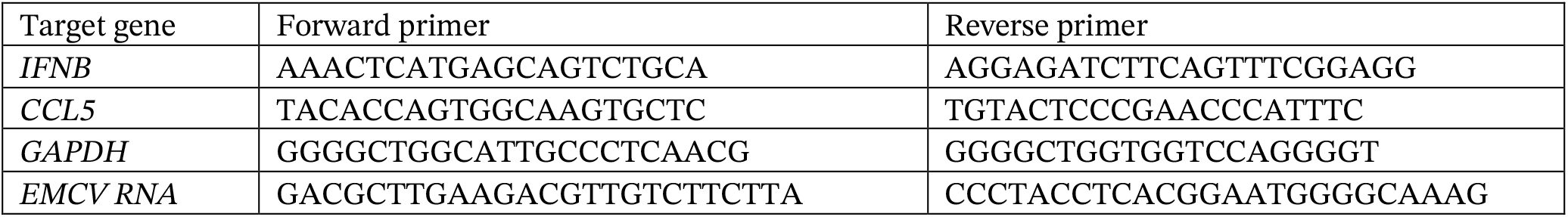
qPCR primer sequences.

**Table S3.** Cryo-EM data collection, refinement, and validation statistics.

|  | #1 apo-MDA5<br>filament<br>(PDB 37NZ) | #2 MA1-bound<br>MDA5 filament<br>(PDB 37NY) | #3 MA2-bound<br>MDA5 filament<br>(PDB 37NW) | #4 MA2-ADP•AIF4-bound<br>MDA5 filament<br>(PDB 37NX) |
| --- | --- | --- | --- | --- |
| **Data collection and processing** |  |  |  |  |
| Magnification | 105,000 | 105,000 | 105,000 | 105,000 |
| Voltage (kV) | 300 | 300 | 300 | 300 |
| Electron exposure (e<sup>-</sup>/Å<sup>2</sup>) | 60 | 60 | 60 | 60 |
| Defocus range (μm) | -0.8 to -1.8 | -0.8 to -1.8 | -0.8 to -1.8 | -0.8 to -1.8 |
| Pixel size (Å) | 0.827 | 0.827 | 0.827 | 0.827 |
| Symmetry imposed | C1 | C1 | C1 | C1 |
| Initial particle images (no.) | 5,433,561 | 5,356,653 | 2,882,609 | 7,945,426 |
| Final particle images (no.) | 169,251 | 610,336 | 348,721 | 378,379 |
| Map resolution (Å) | 3.77 | 3.60 | 2.81 | 3.33 |
| FSC threshold | 0.143 | 0.143 | 0.143 | 0.143 |
| **Refinement** |  |  |  |  |
| Initial model used (PDB) | 4GL2 | 4GL2 | 4GL2 | 7JL2 |
| Map sharpening *B* factor (Å<sup>2</sup>) | -142.9 | -226.4 | -100.4 | -138.5 |
| Model composition |  |  |  |  |
| Non-hydrogen atoms | 11603 | 11572 | 11574 | 11898 |
| Protein residues | 1412 | 1408 | 1406 | 1416 |
| Ligands | 2 | 3 | 3 | 9 |
| R.m.s. deviations |  |  |  |  |
| Bond lengths (Å) | 0.26 | 0.21 | 0.24 | 0.31 |
| Bond angles (°) | 0.44 | 0.38 | 0.41 | 0.57 |
| Validation |  |  |  |  |
| Clash score | 22 | 16 | 9 | 13 |
| Poor rotamers (%) | 1 | 2 | 2 | 2 |
| Ramachandran plot |  |  |  |  |
| Favored (%) | 94 | 94 | 95 | 96 |
| Allowed (%) | 6 | 6 | 5 | 4 |
| Disallowed (%) | 0 | 0 | 0 | 0 |

